# Identification of environmental factors and growth stages in the prediction of fibre yield and fibre quality traits in rain-grown cotton

**DOI:** 10.64898/2026.06.14.732217

**Authors:** Qian Feng, Pierce Rafter, Iain Wilson, Zitong Li, Warren Conaty

## Abstract

**Context:** Understanding how and when environmental conditions influence overall crop performance is crucial for optimising the development of genotypes to a specific breeding target environment. We focused on economically important traits of Australian rain-grown cotton including fibre yield and quality traits, which have not been investigated comprehensively. The aim of the study was to identify relevant environmental factors, and the timing and extent of their impact on rain-grown cotton production.

**Methods:** We used a data driven approach to analyse the relationship between ten climate related environmental factors across various plant growth stages and eight fibre yield and quality traits, using a large-scale field dataset of 9,283 records collected over 23 years at 4 locations, with 53 unique year-location combinations. We applied eight complementary statistical models including stepwise, penalised and Bayesian linear regression, regression-tree based ensemble methods and deep learning frameworks to (1) select the most essential environmental covariates affecting rain-grown cotton production, and (2) evaluate the predictive performance of these models.

**Results:** The environmental impacts on rain-grown cotton production were trait and growth-stage specific. Number of rainy days and solar radiation were identified as the most influential environmental factors for fibre yield traits, vapour pressure deficit at maximum daily temperature was the most influential factor for majority of fibre quality traits. However, each analysed trait was influenced by multiple environmental factors across multiple growth stages (rather than a single factor or a single growth stage). These influential covariates explained a wide range of variation in the traits, accounting for 5.8% to 68.2%. Using the best-fit random forest model, our findings revealed non-linear relationships between key environmental covariates and the traits.

**Conclusions:** Environmental factors at different rain-grown cotton growth stages are key determinants for the performance of end-of-season fibre yield and fibre quality parameters. These findings highlight the need to account for environment conditions when developing cotton varieties optimised for rain-grown production systems. Potential strategies are proposed whereby these key environmental factors can be used to increase the rate of genetic gain in rain-grown cotton production systems.

**Implications:** The results of this study will be crucial for future genetic evaluations and analyses of genotype-by-environment interaction effects in rain-grown cotton, which must account for the influence of the environment on plant performance. Furthermore, these methods can be applied to other species to identify critical growth stages and environmental factors which most influence crop performance.

## 1. Introduction

Over the past 30 years in Australia, between 82% and 99% of the national cotton (*Gossypium hirstum* L.) crop was produced under irrigated conditions (Conaty et al., 2022; Welsh et al., 2022), with the remainder grown under rain-grown conditions where crops rely entirely on stored soil moisture and in-crop rainfall for water. Opportunistic rain-grown cotton in Australia has traditionally been focused in the inland production regions of northern New South Wales and southern Queensland (Figure S1, U.S. Department of Agriculture., 2025). However, interest in rain-grown cotton is increasing as recent studies indicated that large areas currently uncultivated for cotton cultivation could be suitable for its production (Breckwoldt et al., 2014; Rhebergen et al., 2023). Furthermore, expansion of irrigated cotton production in traditional cotton growing regions is constrained by limited water availability, a challenge likely to intensify under climate change (Brodrick and Bange, 2019). In Australia there is a considerable difference in the average lint yield of irrigated cotton (2,600 kg/ha) (Constable and Bange, 2015) and rain-grown cotton (800 kg/ha) (Conaty et al., 2018). Given that the main difference between irrigated and rain-grown cotton production is the quantity and timing of water supply, these, and possibly other, environmental factors have a profound effect on rain-grown cotton yield potential. Even for irrigated production systems, genomic evaluations of cotton indicate that environments (i.e., locations, years) have a large impact on key fibre yield and fibre quality traits (Gapare et al., 2018; Li et al., 2022; Li et al., 2026).

The role of water (rainfall), vapour pressure deficit (VPD), ambient temperature and solar radiation in cotton physiology have been well established through glasshouse experiments and controlled field studies (Conaty et al., 2012; Conaty et al., 2014; Duursma et al., 2013; Monteith, 1977). Cotton plants require water to maintain turgor pressure, transport nutrients, and regulate temperature through evaporative cooling (Blatt et al., 2014; Conaty et al., 2015; Lin et al., 2017). The water requirements of a plant are determined by the rate of evapotranspiration (Hearn, 1979). In the absence of actual evapotranspiration data, VPD, which measures the difference between the atmospheric water vapour pressure and the theoretical atmospheric water vapour pressure at saturation point, may be a good indicator of evapotranspiration (Baker et al., 2007; Castellvi et al., 1996). In addition to water availability and ambient temperature, solar radiation has also been shown to have an impact on cotton physiology (Hake et al., 1991). Single plant analyses have demonstrated that solar radiation is essential for photosynthesis (Reddy et al., 2000) and under normal in-field conditions solar radiation affects plant vigour (Brodrick et al., 2010; Brodrick et al., 2013; Monteith, 1977).

Field studies have noted that total in-season rainfall accounts for only a modest proportion of the variation in lint yield (Blanc et al., 2008; Braunack et al., 2013; Dhaliwal et al., 2022; Traore et al., 2013). Blanc et al. (2008) and Traore et al. (2013) also noted that dry periods were negatively correlated with lint yield, indicating that water stress decreases lint yield. Similarly, Welsh et al. (2022) observed that years classified as hot and dry tended to have the lowest lint yields in rain-grown cotton. Traore et al. (2013) suggested that the distribution of rainfall within the growing season had a greater impact on lint yield rather than the total seasonal rainfall. In agreement with this hypothesis, Dhaliwal et al. (2022) showed that cumulative rainfall between flowering and open boll had a stronger association with lint yield than that from emergence to flowering or over the entire growing season. Previous analyses of rain-grown cotton have also reported associations between lint yield and non-rainfall environmental covariates. Blanc et al. (2008) observed cumulative solar radiation was strongly positively correlated with lint yield. Dhaliwal et al. (2022) noted that after accounting for rainfall, average maximum temperature from flowering to open boll was correlated with lint yield; average maximum temperature between squaring to flowering was also correlated with lint yield, albeit to a lesser extent. In addition to fibre yield, few studies have examined the relationship between environmental factors and fibre quality in rain-grown cotton (Abdelraheem et al., 2020, Ul-Allah et al., 2021, Rehman et al., 2022). Fibre quality refers to key characteristics, such as fibre length, strength, and elongation. Abdelraheem et al. (2020) and Ul-Allah et al. (2021) indicated that water deficit could reduce fibre length and strength, especially during the flowering period. Rehman et al. (2022) found the water stress during the seeding stage may significantly affect fibre micronaire, an indicator of both fibre maturity and fibre fineness.

Together, these glasshouse and field analyses identified key environmental factors influencing fibre yield and fibre quality traits in rain-grown cotton. Therefore, field-based analyses with multiple environment trials (e.g., genomic prediction, genome-wide association studies, or management intervention analyses) need to account for environmental covariates as potential confounding factors. However, it remains unclear which environmental covariates should be included in the analysis. Some recent studies, such as Souaibou et al. (2025), have proposed machine learning approaches to identify key environmental covariates associated with fibre yield and quality traits in irrigated cotton. But to date, no such systematic analysis exists for rain-grown cotton.

From the perspective of statistical methods, identifying key environmental factors and growth stages requires variable selection methods. Accurate prediction of trait performance with relevant predictors requires robust prediction methods. Numerous methods have been proposed in the literature for statistically important variable selection and/or prediction (Grimonprez et al., 2023; He and Wand, 2024; Jarquin et al., 2014; Breiman et al., 2001; Chipman et al., 2010; Schmidhuber, J. 2015, LeCun et al., 2015; Heinze et al., 2018). Conventional methods are generally parametric-based and assume a linear or non-linear relationship between environmental predictors and the target trait. The recent machine learning methods are more flexible since there is no need to specify the exact functional form between predictors and the target, but less interpretable due to implicit model structures. More recently, advanced deep learning techniques have demonstrated superior performance, owning to their layered model structures that enable individual layers to learn sub-domains of complex patterns (Mienye et al., 2024). Despite being largely non-interpretable and often treated as “black boxes”, these deep learning techniques can achieve high prediction accuracy (Molnar, 2020). Without a clear understanding how these methods model the relationships between predictors and the target, identifying the most suitable method for analysing our dataset is challenging. Thus, a systematic evaluation of alternative methods is required to determine the most appropriate approach.

The aim of this study was to identify relevant environmental factors and the timing and extent of their impact on rain-grown cotton production in Australia. To address the aim of this study, we used a large-scale field dataset collected from 23 years at four growing locations in Australia and studied multiple traits (fibre yield and quality) of rain-grown cotton. This was assessed with two classical frequentist methods – SR (stepwise regression) and MLR (multiple linear regression), a penalised method – MLGL (multi-layer group Lasso, Grimonprez et al., 2023), two Bayesian methods – BGAM (Bayesian generalized additive model, He and Wand, 2024) and LMM (linear mixed model, Jarquin et al., 2014), two tree-based ensemble methods - RF (random forest, Breiman et al., 2001) and BART (Bayesian additive regression tree, Chipman et al., 2010), as well as a deep learning method – DNN (deep neural network, Schmidhuber, J. 2015, LeCun et al., 2015). These methods were used to identify the influential environmental factors/growth stages and predict trait performance. Our analysis contributes to the small body of literature on the association between environmental covariates and traits’ performance in rain-grown cotton and are important as they can be applied to cotton breeding programs aiming to produce varieties where productivity is optimised to rain-gown production systems, setting a foundation for further genomic analysis of rain-grown cotton (including more accurate estimation of breeding values). While the results of this study are only directly applicable to the Australian rain-grown cotton production context, through method comparison we also aim to identify and recommend the most informative statistical method for the translation of this research into other breeding contexts.

## 2. Materials and methods

### 2.1. Trait data

Phenotype records for lint yield (LY), lint percentage (LP), and six different fibre quality traits including fibre length (LEN), strength (STR), micronaire (MIC), elongation (EL), short fibre index (SFI) and uniformity (UNI) (see detailed trait definitions in Li et al., 2022) were available for 4,316 genetically distinct cotton breeding lines grown under rain-grown conditions over a 23-year period from 1999 to 2021 at four separate locations in northern New South Wales and southern Queensland, Australia. In total there were 53 distinct year-location combinations. Half (54.4%) of breeding lines studied were grown in a single site-year and the remaining (45.6%) breeding lines were grown in more than one site-year. On average, each of the breeding lines were grown in more than two different site-year, so that in total 9,283 trait records were available for the present study. The four growing locations were Bellata (29°55’ S, 149°47’ E), Dalby (27°10’ S, 151°15’ E), Narrabri (30°12’ S, 149°36’ E), and North Star (28°57’ S, 150°25’ E). At every growing site the soil was a uniform grey cracking clay (USDA soil taxonomy: Typic Haplustert; Australian soil taxonomy: Grey Vertosol) with clay contents of approximately 65% in Bellata, 70% in Dalby, 60% in Narrabri, and 55% in North Star (Isbell, 1996). Due to seasonal nature of water availability, rain-grown cotton was not grown at every location in each year (**Table 1**).

**Table 1.** The growing seasons and number of phenotype records for each of the four cotton growing sites: Bellata (29°55’ S, 149°47’ E), Dalby (27°10’ S, 151°15’ E), Narrabri (30°45’ S, 150°34’ E), and North Star (28°57’ S, 150°25’ E).

| Site | Growing season | Number of records |
| --- | --- | --- |
| Bellata | 2006-2009, 2011, 2013, 2015-2017, 2021 | 1,823 |
| Dalby | 1999-2001, 2003-2005, 2007-2011, 2015-2018, 2020-2021 | 2,059 |
| Narrabri | 1999-2017, 2019-2021 | 4,936 |
| North Star | 2001-2004 | 465 |

All phenotype data was collected from experiments conducted as part of the rain-grown component of the CSIRO Cotton Breeding program. The Narrabri site was located at the program’s core research base at the Australian Cotton Research Institute (ACRI), while all other sites were located on commercial rain-grown cotton enterprises. Experiments were laid out in row-column designs with four replicates, generated from CycDesigN software (VSN International, Hemel Hempstead, UK). Each plot consisted of two 14-16 m rows of cotton (depending on the individual experiment). A row spacing of 1 m was used with a planting density of about 8-10 plants m^−2^, where depending on the enterprise’s standard practices either a single skip row or double skip row configuration was used (i.e., two rows in, 1 row out or two rows in, 2 rows out, respectively). Therefore, each plot consisted of two rows of cotton, with either one or two adjacent empty rows on each side of the plot. Management for all field experiments followed then or current commercial practices for Australian rain-grown (i.e., dryland) cotton production (Cotton Research and Development Corporation, 2025). Each experiment was managed according to its individual requirements for weed and pest control, with all plots receiving the same management regime.

### 2.2. Environmental factors

The environmental data consisted of daily records for solar radiation (MJ/m^2^), maximum and minimum temperature (℃), rainfall (mm), and relative humidity at maximum and minimum temperature (%). These daily records were available for the entire growing season, from planting to harvest. Environmental data were collected from weather stations (OzForecast, Narrabri Australia) located close to the cotton growing sites. When weather station data were not available, environmental data were obtained from the SILO climate database (Jeffrey et al., 2001). Weather stations were not always kept in a fixed location from year to year, as such there was some variance in the distance between the growing site and the nearest weather station. The distances between the site location and the nearest weather station were between approximately: 2.2km and 5.2km for Bellata, 0.1 km and 6.2 km for Dalby, 0.8 km and 1.4 km for Narrabri, and was consistently 2.9 km for North Star.

Water vapour pressure at the saturation point for daily maximum temperature was calculated in kilopascals (kPa) using the Tetens formula (Xu et al., 2012):

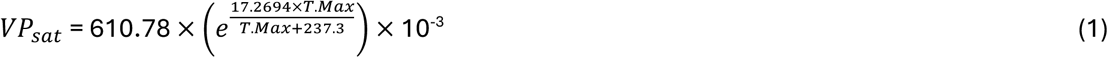

where T. Max represents maximum daily temperature in Celsius and VP_sat_ is the atmospheric water vapour pressure at the saturation point. Daily VPD_max_ was calculated using the following formula (Brun et al., 2022):

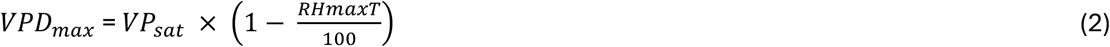

where RHmaxT is the relative humidity at maximum daily temperature expressed as a proportion of maximum humidity and VP_sat_ is as described in **Eq. 1**. A high VPD_max_ indicates hot and dry environment while low VPD_max_ indicates cold and wet environment. VPD_max_ was added as one of the environmental factors analysed in the present study.

To account for the extreme climate events including heatwaves and drought spells, three additional environmental factors were considered, number of cold days (< 12℃), number of hot days (> 35℃), and number of rainy days during each growth stage (see section below for the definition of growth stage).

### 2.3. Cotton growth stages

Although the cotton plant is indeterminant (Snider et al., 2021), when cotton is grown as an annual crop its development can be considered in terms of seven distinct phenological growth stages (Bange et al., 2022; Constable and Shaw, 1988). The phenological growth stages are planting to emergence, emergence to first floral bud, first floral bud to first flower, first flower to peak flowering, peak flowering to open boll, open boll to maturity, and maturity to harvest. Growth-stage development in cotton can be approximated as a mathematical function of day-degrees (Constable and Shaw, 1988). The day-degree is a thermal-time metric intended to reflect the potential rate of plant development based on the temperature range for a given day. Day-degrees were calculated using the following phenology model (Bange et al., 2022; Constable and Shaw, 1988):

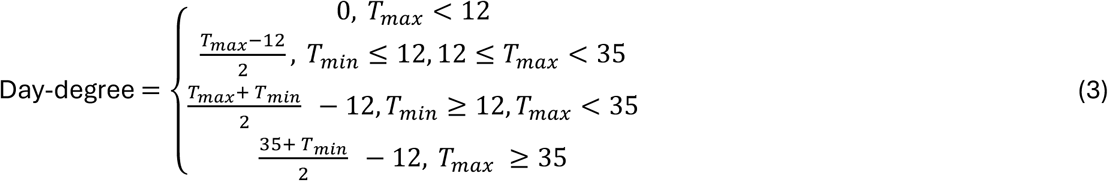

where T_max_ and T_min_ are the maximum and minimum temperatures in Celsius recorded for a given day. The start and end cumulative day-degrees assumed to coincide with the phenological growth-stages of rain-grown cotton are given in **Table 2**. The developmental phase of the plant is assumed to last for the duration of the growth stage(s), for example flowers are expected to remain in bloom for the duration of the growth-stages from first flower to the end of peak flowering.

**Table 2.** The cumulative day-degrees for the seven distinct growth stages of cotton as defined by the physiological development of the cotton plant. The seven growth stages refer to (1) from planting to emergence, (2) from emergence to first square, (3) from first square to first flower, (4) from first flower to peak flower, (5) from peak flower to open boll, (6) from open boll to maturity, (7) from maturity to harvest.

| Growth stage | Cumulative day-degree start | Cumulative day-degrees end | Phenological feature |
| --- | --- | --- | --- |
| One | 0 | 80 | Emergence |
| Two | 80 | 505 | First square |
| Three | 505 | 777 | First flower |
| Four | 777 | 1065 | Peak flower |
| Five | 1065 | 1352 | Open boll |
| Six | 1352 | 1815 | Maturity |
| Seven | 1815 | NA | Harvest |

*2.4. Environmental data preprocessing*

### 2.4. Environmental data preprocessing

The physiological requirements of cotton plant are likely to vary across different developmental stages (Constable and Shaw, 1988; Snider et al., 2021). Therefore, the influence of environmental conditions on the plant hypothetically also depends on the stage of plant development. We therefore counted the number of cold/hot/rainy days within each growth stage. While for the remaining seven daily-based environmental factors, we processed them with two different approaches.

1. We grouped daily environmental records into growth-stage based records. For solar radiation, we followed previous work (Blanc et al., 2008) and calculated the cumulative solar radiation within each growth stage. For the remaining factors, the average of daily records within each growth stage were calculated.
2. We modelled the temporal trends of environmental factors using the B-spline regression (Eilers and Marx, 1996), and the estimated B-spline regression coefficients were used as environmental covariates (Savino and Lévy-Leduc, 2025).

In above scenario 1, a total of 70 environmental covariates (10 environmental factors at 7 growth stages) were available. Each environmental covariate (EC) is the combination of an environmental factor (like rainfall, solar radiation) and a plant growth stage. In scenario 2, eight regression coefficients per daily environmental factor were generated (see more in supplementary text), and this resulted in 77 ECs in total (56 ECs from daily factors, 21 ECs from the number of cold/hot/rainy days across growth stages). Although two scenarios corresponded to slightly different number of ECs (70 vs. 77), the same workflow for statistical analyses was applied to both. For brevity, we illustrate using the 70-EC scenario below, the same mechanisms apply to 77-EC scenario.

These environment data were finally matched to the trait data, based on year and location. The resulting dataset consisted of 9,283 trait records and 70 ECs. Across the entire dataset, the summary statistics (mean, standard deviation, minimum-maximum range) for all analysed traits/ECs and related abbreviations are in **Table 3**.

**Table 3.**
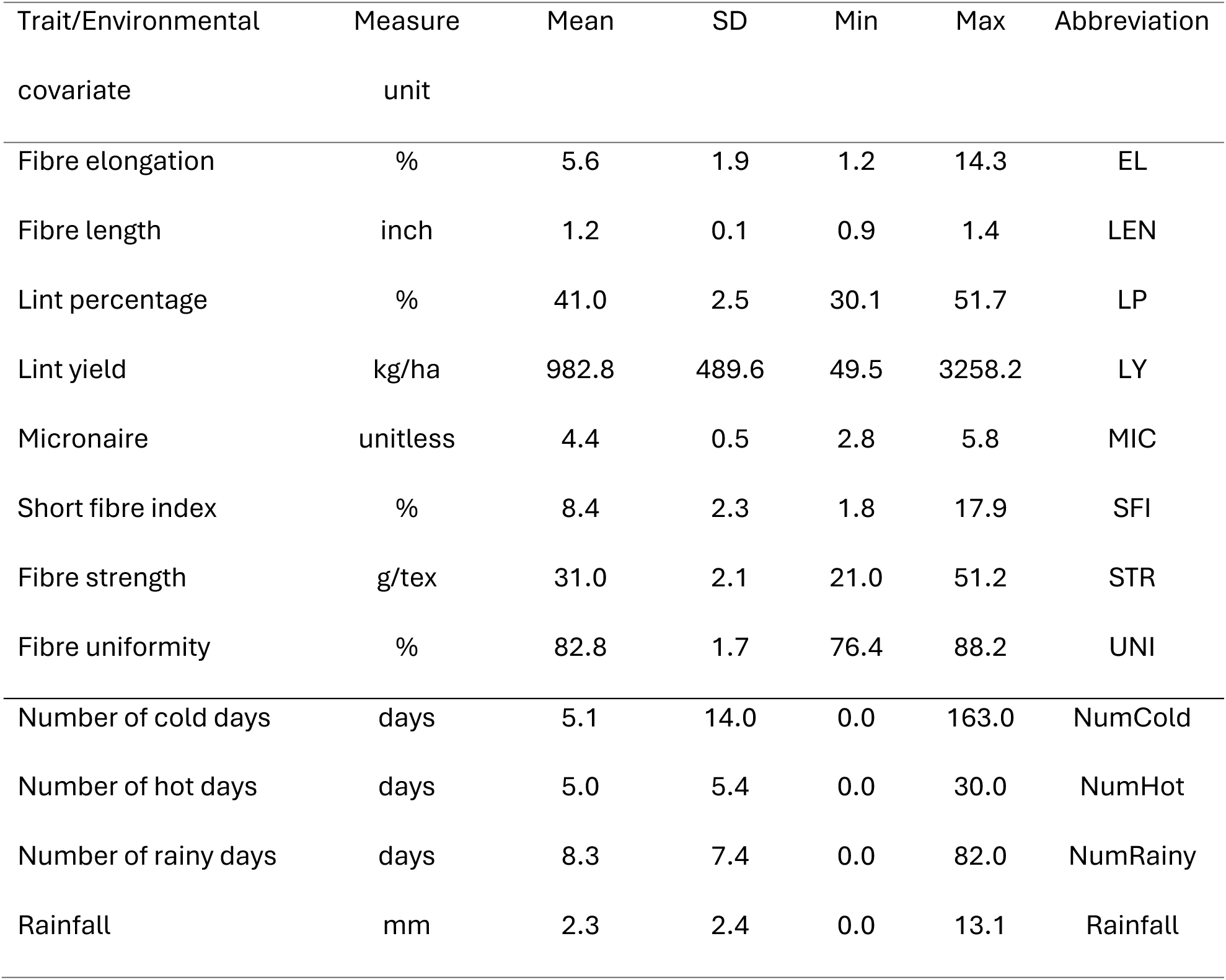
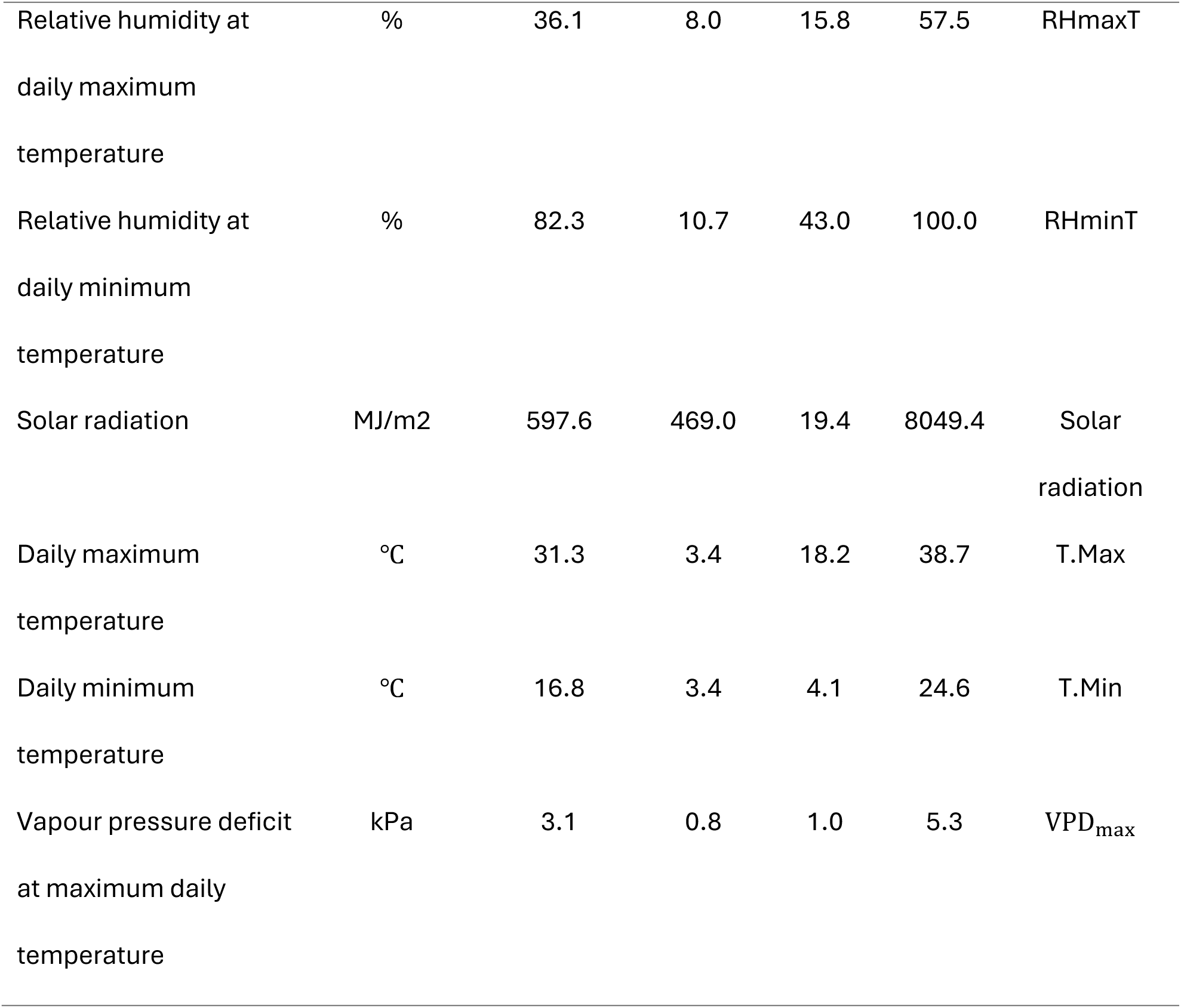
The descriptive statistics (mean, standard deviation, minimum and maximum value), measure unit and abbreviation of eight fibre yield and fibre quality traits of rain-grown cotton, ten environmental factors (per year/location/growth stage) from the large-scale field dataset collected over 23 years at 4 growing locations in Australia.

### 2.5. Workflow for statistical analyses

A workflow utilising statistics and machine learning approaches was used to identify the environmental factors that influence fibre yield and fibre quality traits in rain-grown cotton production, when these effects occur and how they manifest (**Figure 1**). The workflow consists of three steps:

1. Collect and preprocess the trait and environmental data. We computed and normalised each EC.
2. Determination of the most essential ECs. Six different variable selection methods were applied to the same data. A consensus strategy was adopted where a variable (i.e., EC) was selected if it was selected by at least four out of six methods.
3. Study the manifestation of key ECs on the trait.

- We determined the best performing model (most accurate in the prediction) by applying seven different prediction methods. Cross validation was used to evaluate methods’ predictive performance.
- With the optimal prediction model, we inferred the importance of influential ECs (from Step 2) and determined the type of association (e.g., linear, non-linear, positive, negative, etc.)

**Figure 1.**
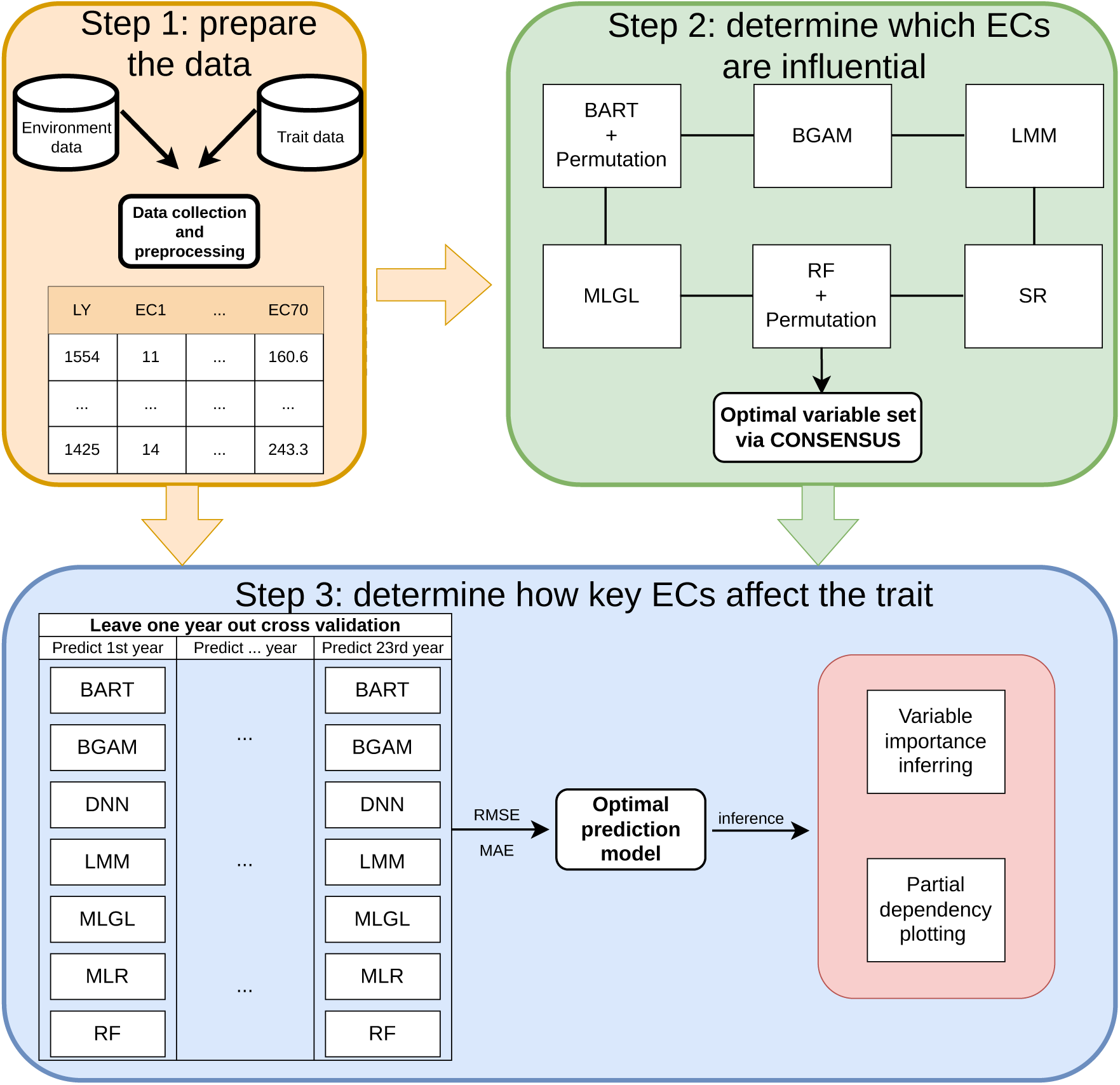
A graphical representation of proposed workflow to identify the most essential environmental factors, their timing and specific impacts on fibre yield and quality traits of Australian rain-grown cotton. We started with data collection and preprocess. To identify influential environmental covariates (ECs), six complementary variable selection methods were used separately, with optimal set of ECs chosen by the consensus. Next, the most accurate prediction model was determined after comparing the performance of seven different prediction models. The final model inference was conducted with the optimal prediction model for specific environmental impacts. See Abbreviations for the full names of methods used.

We applied the same procedure to separately analyse the eight traits.

#### 2.5.1. Statistical methods introduction

In this study, each of the eight traits analysed was modelled separately as:

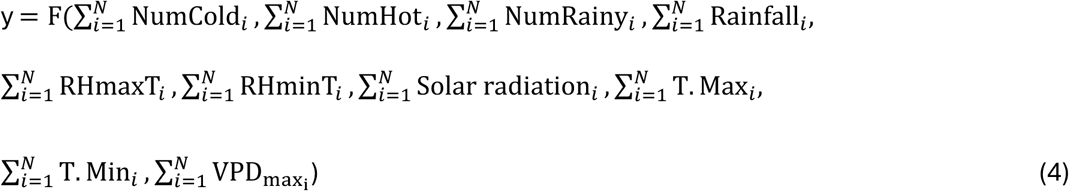

where y is the trait of interest and N (=7) represents the number of growth stages. The function *F*(·) represents a general function form describing the relation between ECs and the trait. Different variable selection and prediction methods correspond to different specifications of *F*(·), including differences in model structure, assumptions about linearity, and parameter estimation. As such, each method defines a specific form of *F*(·), while the underlying set of 70 ECs remains consistent across methods.

Two classical methods were used as separate baselines: bidirectional stepwise regression (SR) for variable selection and multiple linear regression (MLR) for prediction. The MLR is the foundational model in prediction and includes all user-specified predictors, while SR automates MLR process in each step to either include or exclude variables. The MLR model used was as below

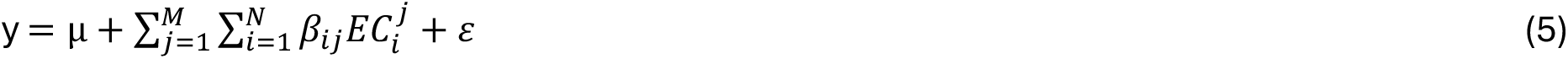

y is the trait of interest, µ is the intercept, and 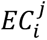 refers to the *j*^*th*^ environmental factor at *i*^*th*^ growth-stage. *M* is the total number of environmental factors analysed, *β*_*ij*_ denotes its effect size, and ε is the independent and identically distributed error term. We implemented MLR by R function “lm” and parameters were estimated using well-known ordinary least square (OLS). SR was implemented via R function “step” with optimal model determined by Bayesian Information Criteria.

MLGL (Grimonprez et al., 2023) is a penalised regression methods which can do both variable selection and prediction. The MLGL method was designed for variable selection when variables exhibit correlation and/or grouping structure. It consists of three major steps, firstly, a hierarchical clustering of predictors is performed to generate a hierarchical structure of nested groups of predictor variables. Group-Lasso (Yuan and Lin, 2006) is applied to the variable groups derived from the hierarchical clustering step, where entire groups are penalised and selected by group-wise *l*_1_ norm penalty. Finally, a hierarchical multiple testing is used to control the false discovery rate (FDR) among the selected groups. In Group-Lasso, on top of the OLS, a penalty term was added:

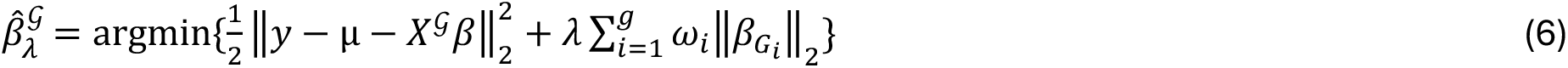

where *G* is the union of all groups across all hierarchy levels, 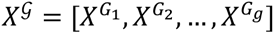 is a design matrix handling all partitions from all hierarches *G* = {*G*_1_, *G*_2_, …, *G*_g_}, *λ* > 0 is the regularization parameter and *ω*_*i*_ is a weight associated with group *G*_*i*_. This penalty could shrink coefficients of groups to zero for performing variable selection and avoids overfitting. The MLGL method was implemented using the “MLGL” R package.

BGAM (He and Wand, 2024) is a Bayesian non-linear regression method that assumes the effects of ECs to be zero, linear or non-linear, and employs specified priors to shrink regression parameters toward these alternatives, thereby facilitating both variable selection, flexible effect estimation, and prediction. In BGAM, each trait in the present study was modelled as a generalized additive model:

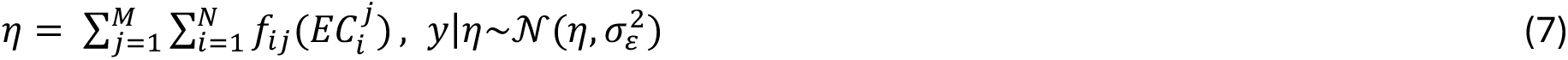

where *f*_*ij*_(*x*) is the smooth function for the EC, 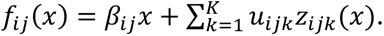 Here, *β*_*ij*_ represents the linear effects, and *u*_*ijk*_ are the coefficients associated with the spline basis function *z*_*ijk*_(*x*), capturing the non-linear component of the effect. The spline basis *z*_*ijk*_(*x*) are constructed to be orthogonal to the linear component (Chouldechova and Hastie, 2015), ensuring identifiability and enabling a clear separation between linear and non-linear effects. The Markov Chain Monte Carlo (MCMC) are used to draw samples to approximate the posterior distributions of parameters, and these samples are then used for model inference. This method was implemented with the R package “gamselBayes”. We first generated 5,000 samples as burn-in and discarded them. Then, 20,000 samples were drawn and thinned by retaining every 10th sample to reduce serial correlation. This resulted in 2,000 samples to approximate the posterior distributions.

The linear mixed model (LMM) (Jarquin et al. 2014) used for both variable selection and prediction in this study is defined as

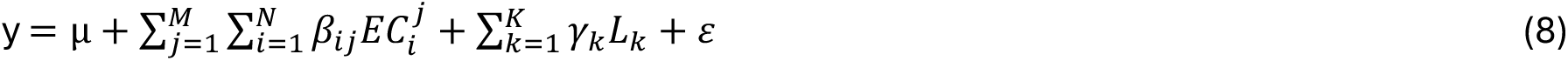

where *L*_*k*_ denotes the design matrix for *k*^*th*^ cotton line, *K* is the total number of cotton lines in the data, and *γ*_*k*_ is the effect size of each cotton line. Cotton lines were treated as random effects and ECs as fixed effects. Thus, each trait was modelled by the overall mean (µ), ECs and cotton line effects, and the error term (*ε*). In the implementation, *β*_*ij*_ was assigned a spike and slab prior (BayesC, Haber et al., 2011), assuming that only a proportion of the effects are non-zero, while γ_*k*_ was assigned with a Gaussian prior. All parameters were estimated using MCMC. This model was implemented using “BGLR” R package (Perez and de los Campos, 2014), and the number of MCMC samples, burn-in, thinning intervals were the same as those used in BGAM.

RF and BART are ensemble methods which either combine or average the results of many decision tree models into a single overall ensemble model (Breiman, 2001; Chipman et al., 2010). In RF, each decision tree is trained independently on a bootstrap sample of the data. The performance of each decision tree is evaluated using the out-of-bag observations (those not included in the bootstrap sample). The final prediction of the RF is obtained by aggregating the predictions of all trees (Breiman, 2001). The RF model was implemented with the “randomForest” R package (Liaw and Wiener, 2002).

A two-stage grid search approach (Ramadhan et al., 2017; Sumathi, 2020) was used to determine the optimal set of user-defined hyperparameters, i.e., the number of trees in the model *ntree*, the number of independent variables randomly selected for each tree *mtry*, and the minimum number of observations used for tree leaves *nodesize*. Given the recommended *ntree*, *mtry*, *nodesize* were 500, 23, 5 separately from “randomForest”, we set the range of each hyperparameter as *ntree* ∈ {250, 500, 750, 1000}, *mtry* ∈ {10, 20, 30, 40}, *nodesize* ∈ {5, 10, 15, 20} in the first coarse stage. The second stage further multiples each of the best parameter chosen from the first stage by {0.7, 0.8, 0.9, 1.0, 1.1, 1.2, 1.3}, allowing a finer search. The optimal parameters in each stage were those producing the smallest mean square error.

Unlike RF, which builds many independent trees and aggregates their predictions, the BART method represents the response as a sum of multiple trees. It adopts a fully Bayesian framework by placing priors on tree structures and terminology parameters, then employs MCMC to iteratively sample from the posterior distribution of these trees. This approach allows BART to capture complex nonlinear relationships while providing coherent uncertainty estimates for predictions (Chipman et al., 2010; Hill et al., 2019). The BART method was evaluated using the “BART” R package (Sparapani et al., 2021). The default parameters for the BART which included 1,000 MCMC samples with a burn-in of 100 were used because previous studies on a diverse set of datasets (Chipman et al., 2010) and cotton (Li et al., 2022) have demonstrated robust results using the default parameters recommended by the developers of the BART method. Moreover, the MCMC convergence for each trait was diagnosed and passed by “coda” R package (Plummer et al., 2006).

Although both RF and BART are powerful predictive methods, neither of them can perform explicit variable selection. Instead, we used the variable importance values generated from each method combined with a permutation test (Dudoit and van der Laan, 2008) to select significant variables. Specifically, the phenotype data for a given trait were randomly reshuffled 1,000 times and each permuted dataset was analysed by either RF or BART. This procedure generated an empirical null distribution of variable importance scores for each EC, which can then be used to access the statistical significance (*P* < 0.05) of observed importance values in the original data. For RF, the variable importance was measured by the mean decrease in accuracy after excluding the variable in question. For BART, it was measured by the average count of the variable used for growing trees among MCMC samples.

To determine the key influential ECs, we implemented a consensus variable selection strategy, where only ECs identified with at least four out of six methods were retained. The aim with this approach was to reduce the chance of inducing false positives, albeit at the potential cost of excluding some true positives, thereby prioritizing robustness over sensitivity. To further measure the proportion of variance explained by these key ECs per trait, the adjusted *R*^2^ from multiple linear regression was calculated.

We also assessed a deep neural network (DNN) model for prediction, which is particularly effective at capturing complex and non-linear relationship among data (Schmidhuber, J. 2015, LeCun et al., 2015). The architecture of our DNN model was a fully connected feed-forward design, consisting of an input layer (with 70 neurons, defined by number of ECs), two hidden layers (with 64 and 32 neurons) and an output layer (with single neuron). Neurons were fully connected at adjacent layers, enabling dense information flow throughout the network. The ReLU activation function was employed at each hidden layer. Our model was trained using the Adam optimization algorithm with mean square error as the loss function. Training was performed for 100 epochs with a batch size of 32, and 20% of the training data was held out for validation. After training, the fitted model was used to generate predictions on the independent test dataset. This model was implemented using R package “keras3” (Kalinowski et al., 2025).

#### 2.5.2. Predictive accuracy assessment

All ECs, instead of only selected ECs from step 2, were used as predictors for selecting the best prediction model, this avoids data being used twice for first doing variable selection and then prediction (which will over-estimate prediction power). To evaluate predictive performance of a model on untested data, a leave-one-year-out cross-validation approach was used to compute the predictive accuracy. A single year of data was used as the validation dataset and the remaining data from all other years were used as the training dataset. We chose the leave-one-year-out rather than leave-one-site-out cross-validation to avoid the data leakage due to the geographically close growing sites. Since the dataset spanned 23 years (1999-2021), the validation approach process was repeated 23 times such that each year of data had predicted and observed trait values. We evaluated the predictive accuracy by the root mean square error (RMSE) and mean absolute error (MAE) using the observed and predicted values for each test year.

#### 2.5.3. Model inference

Once the optimal prediction model was determined, we calculated the partial dependence by predicting the trait using the trained model on a dataset where all variables were fixed at their mean values, except for the EC of interest, which was varied across its range.

## 3. Results

In the following, we describe the results using ECs from scenario 1 below, please refer to supplementary materials for scenario 2 (supplementary text, Tables S1-S2, Figures S4-7), since we observed very similar results between two scenarios, including both variable selection and prediction performance.

### 3.1. Variable selection analysis

The number of identified influential ECs across six variable selection methods were diverse (**Table 4)**. Among all method used, ensemble-based methods selected out the smallest set of variables, and the baseline stepwise method generally returned the biggest set. Specifically, BART was the most conserved in selecting variables, followed by RF. In contrast, SR selected the greatest number of variables for five out of eight traits analysed.

**Table 4.** The number of identified influential environmental covariates (ECs) using 6 variable selection methods and consensus strategy for fibre yield and fibre quality traits of Australian rain-grown cotton. The consensus set was defined by the ECs identified ≥ 4 variable selection methods. See Abbreviations for the full name of each method.

| Trait | BART +<br>Permutation | BGAM | LMM | MLGL | RF +<br>Permutation | SR | Consensus |
| --- | --- | --- | --- | --- | --- | --- | --- |
| EL | 5 | 31 | 45 | 0 | 15 | 50 | 4 |
| LEN | 4 | 26 | 45 | 70 | 17 | 48 | 14 |
| LP | 1 | 8 | 37 | 47 | 25 | 47 | 8 |
| LY | 9 | 39 | 47 | 70 | 12 | 50 | 24 |
| MIC | 3 | 11 | 43 | 15 | 11 | 49 | 2 |
| SFI | 1 | 12 | 41 | 47 | 40 | 51 | 18 |
| STR | 0 | 18 | 48 | 47 | 28 | 45 | 13 |
| UNI | 1 | 27 | 46 | 47 | 16 | 51 | 19 |

With consensus strategy, the final set of influential ECs identified per trait are visualised in **Figure 2**. LY was affected by the greatest number of influential ECs (24 out of 70), and the variance explained was up to 68.2%. Half of the fibre quality traits (LEN, SFI, UNI) were affected by a slightly smaller number of ECs and variance explained ranged from 40.6% to 53.7%. For the remaining fibre quality traits (EL, MIC and STR), only a few ECs were selected out and the proportion of variance explained were below 20%. The relationship between number of influential ECs and proportion of variance explained appeared to be linear (*P* = 0.005, Figure S2).

**Figure 2.**
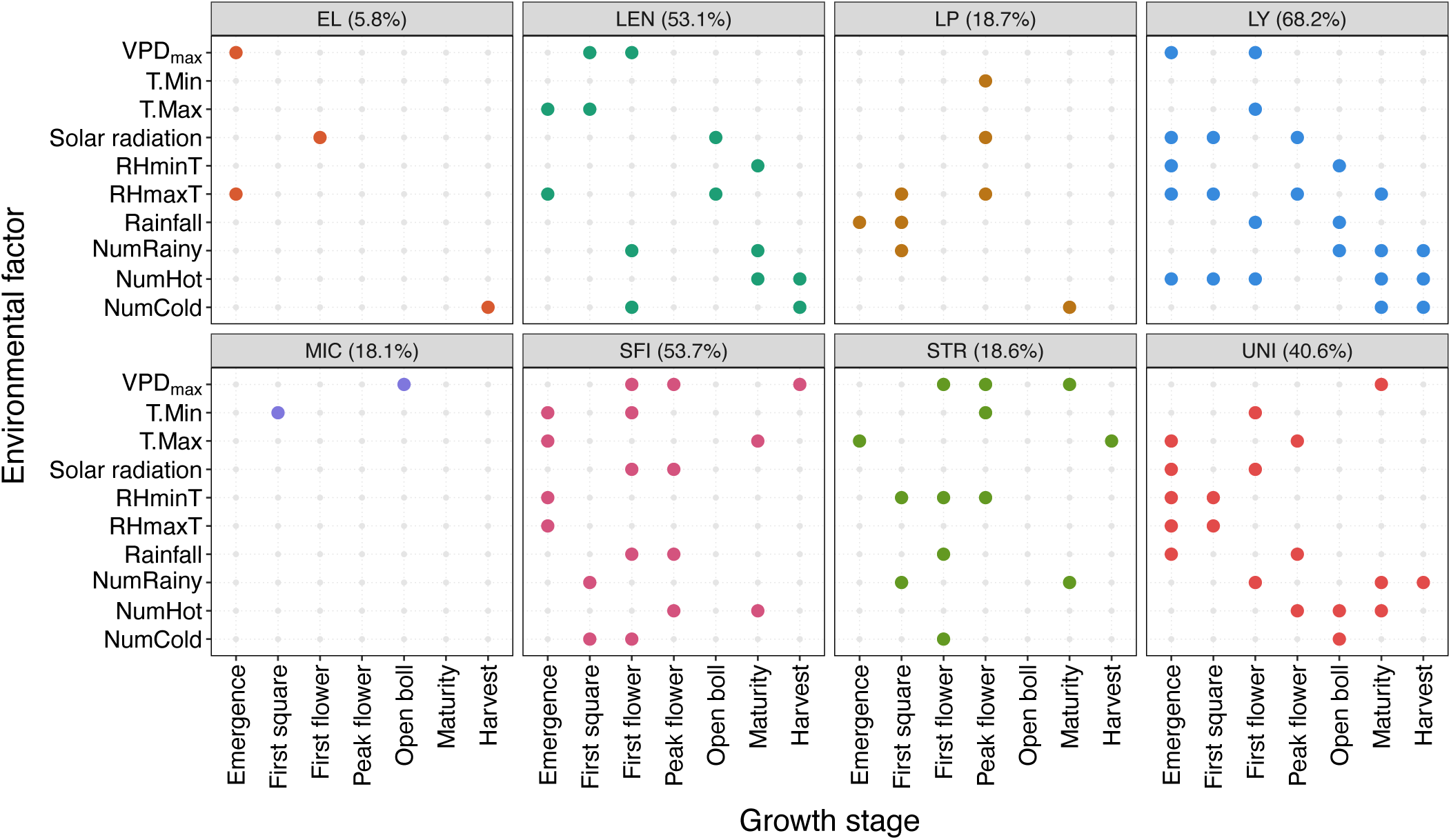
The identified influential environmental factors and growth stages for fibre yield and fibre quality traits of Australian rain-grown cotton. Eight panels represent eight traits separately, the rows and columns in each panel represent various environmental factors and growth stages. If an environmental factor at a growth stage was found to influence a trait, a coloured dot in the corresponding panel was highlighted. The number in the bracket denotes the proportion of variance explained by the identified influential environmental covariates. EL, LEN, LP, LY, MIC, SFI, STR and UNI stand for fibre elongation, length, percentage, yield, micronaire, short fibre index, strength and uniformity separately.

The analyses revealed that the influential ECs were trait specific. Despite this, across all environmental factors analysed, each factor appeared to be influential for at least half of traits, although the specific influential growth stage varied by trait. Likewise, at each growth stage, there were always environmental factors influencing at least half of the traits analysed, although the specific factors differed among traits.

### 3.2. Model prediction accuracy

For each prediction model, the average RMSE and standard deviation calculated with the observed and predicted trait values across years was given in **Figure 3**. Results indicate RF was the most accurate prediction model for seven out of eight traits analysed. Despite BART was the most accurate model for SFI, we found the RMSE values were very similar between RF and BART. In fact, the RMSE values were comparable between these two ensemble models across all traits analysed, the performance of penalised regression (MLGL) ranked the second, and the baseline MLR method performed the worst. We also drew similar conclusions when MAE was used as the criterion to measure prediction accuracy (Figure S3).

**Figure 3.**
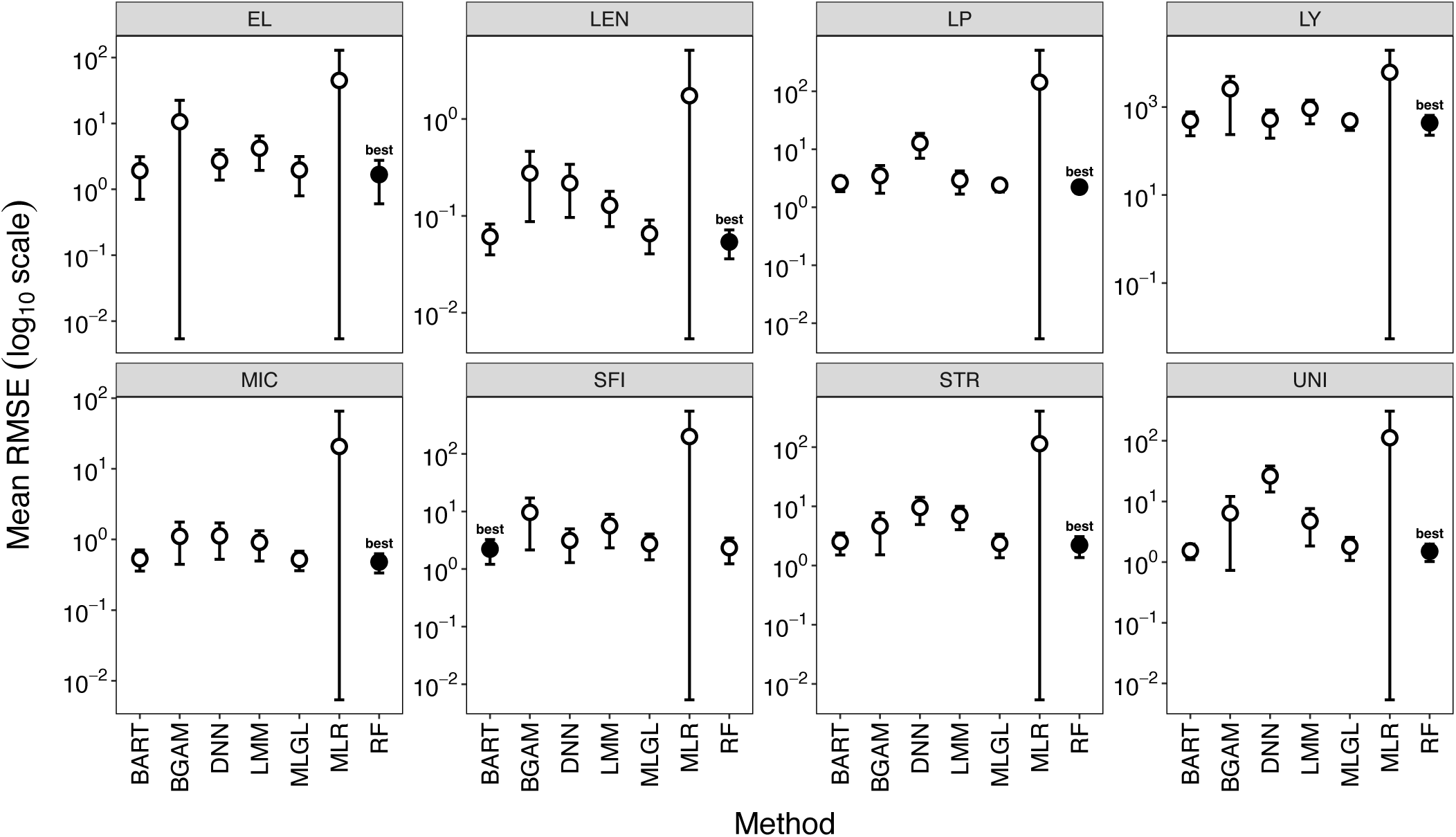
Prediction accuracy measured by root mean square error (RMSE) of seven different methods for predicting fibre yield and fibre quality traits of Australian rain-grown cotton. Each error bar refers to the Mean ± SD RMSE across years, and filled circle denotes the best method (smallest RMSE). BART, BGAM, DNN, LMM, MLGL, MLR and RF represent the Bayesian Additive Regression Tree, Bayesian Generalized Additive Model, Deep Neural Network, Linear Mixed Model, Multi-Layer Group Lasso, Multiple Linear Regression, Random Forest.

Scatter plots of mean trait values for each year-location against the predicted values from RF model for all analysed traits were shown in **Figure 4**. For most traits (except LY and STR), the 95% confidence intervals of the linear regression between predicted and observed trait values include the 45% reference line (where predicted equals observed), indicating a good predictive performance of RF model. Therefore, we chose RF as the default approach for subsequent model inference.

**Figure 4.**
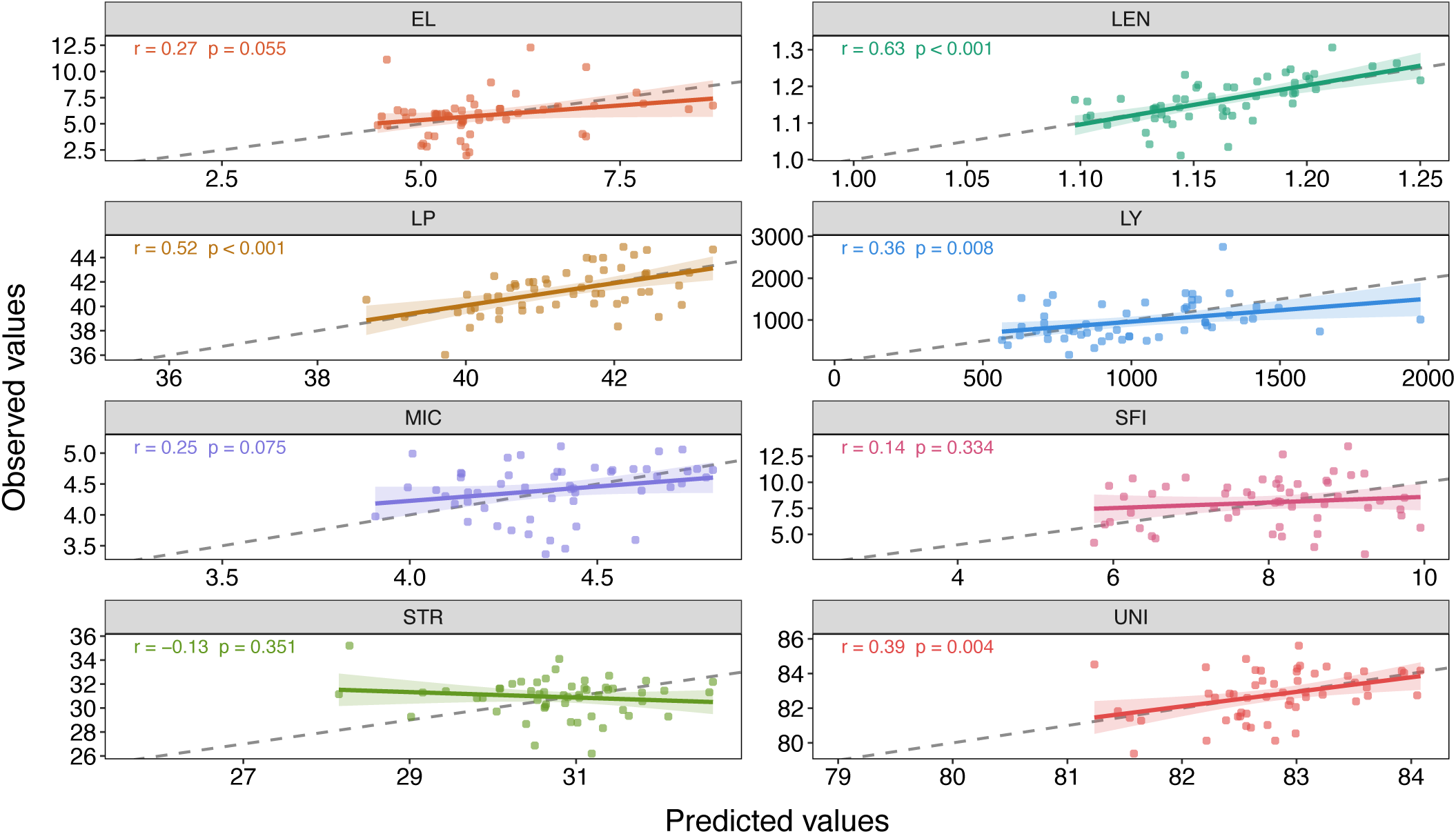
Observed versus predicted mean trait values per year-location from the best-fit random forest model. Each panel represents a trait. Each dot in the panel represents a year-location combination. The dashed line is a 45% reference line, and the solid line is a linear relationship between the observed and predicted mean traits values. The shadow area denotes the 95% confidence interval. The correlation *r* between predicted and observed values and *p* value for the fitted regression relationship were reported in the top left of each panel.

### 3.3. Variable importance and partial dependency

The importance of influential ECs was assessed by the *P* values from permutation test based on the best-fit RF model (from Step 2). **Figure 5** illustrates the importance of influential ECs for various fibre yield and fibre quality traits (ECs with *P* > 0.05 are not shown). Overall, multiple environmental factors across multiple growth stages appeared to influence each trait analysed; no trait was solely influenced by a single environmental factor or a single growth stage.

**Figure 5.**
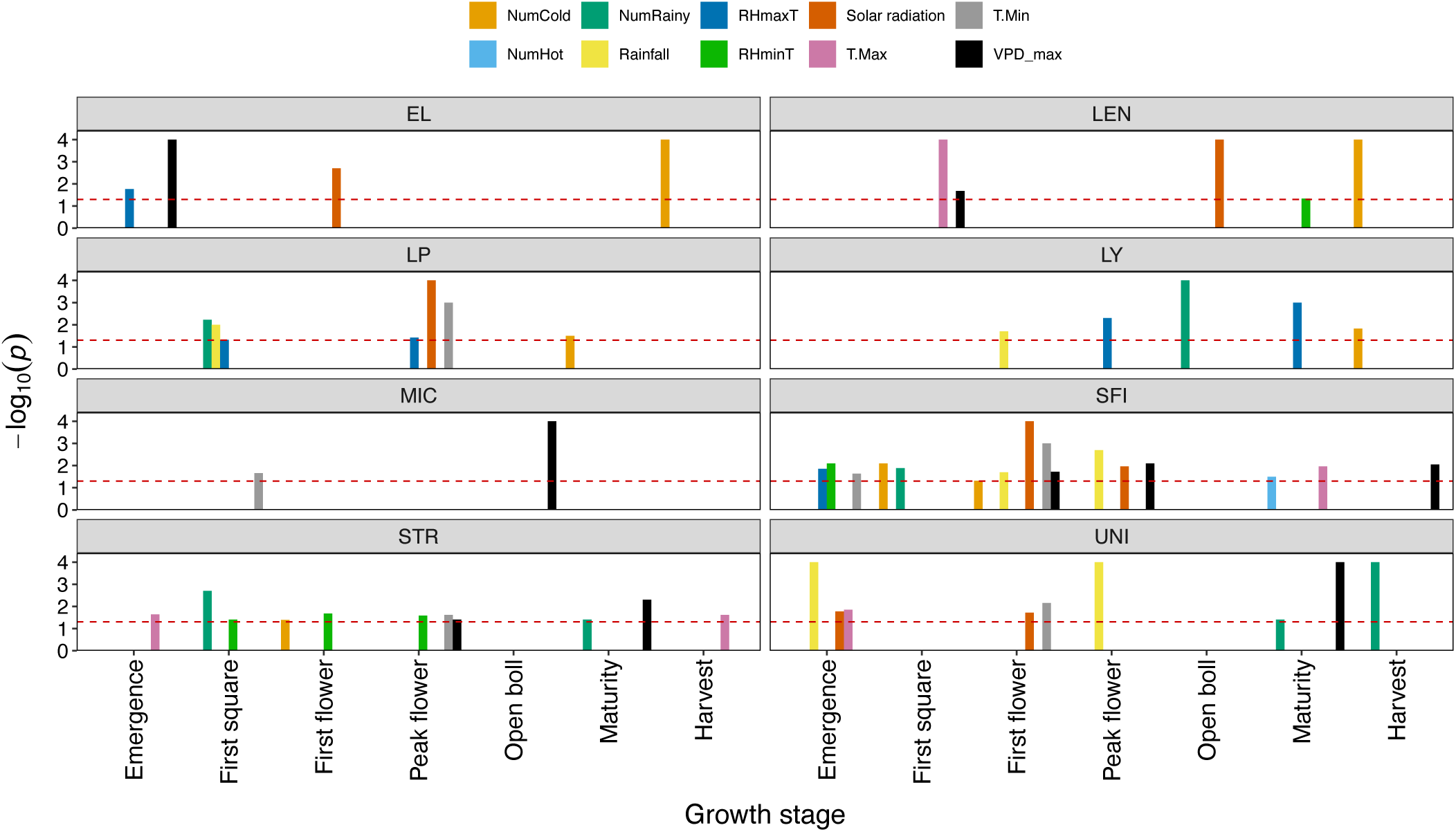
The importance of influential environmental covariates (ECs) for fibre yield and fibre quality traits of Australian rain-grown cotton. Each panel in this graph represents a trait. The importance values of ECs were computed by the *p* value of permutation test using the best-fit random forest model. The *p* value was −*log*_10_ transformed for better visualisation. The importance of ECs was represented by the height of coloured bars which refer to different environmental factors. ECs with a reported *p* = 0 were capped at -*log*_10_(0.0001) = 4, shown at the top. The horizontal line refers to 5% significance threshold. The full name of each trait and environmental covariate abbreviation is shown in **Table 3**.

The number of rainy days, cumulative solar radiation and mean VPD_max_ were identified as the most important environmental factors, each significantly influencing at least five of the eight traits analysed. Specifically, the number of rainy days significantly affect two fibre yield traits (LY and LP), and half of fibre quality traits (SFI, STR and UNI). The relationship between this factor and the traits depended on the phenological growth stage. In particular, LY and STR are two economically important traits in cotton. The number of rainy days from peak flowering to open boll period significantly influenced LY, while the number of rainy days from the plant’s emergence to first square and from open boll to maturity stage significantly influenced STR. Moreover, the number of rainy days within a certain range at the most influential growth stage showed a positive relationship with both LY and STR (**Figure 6a,d**).

**Figure 6.**
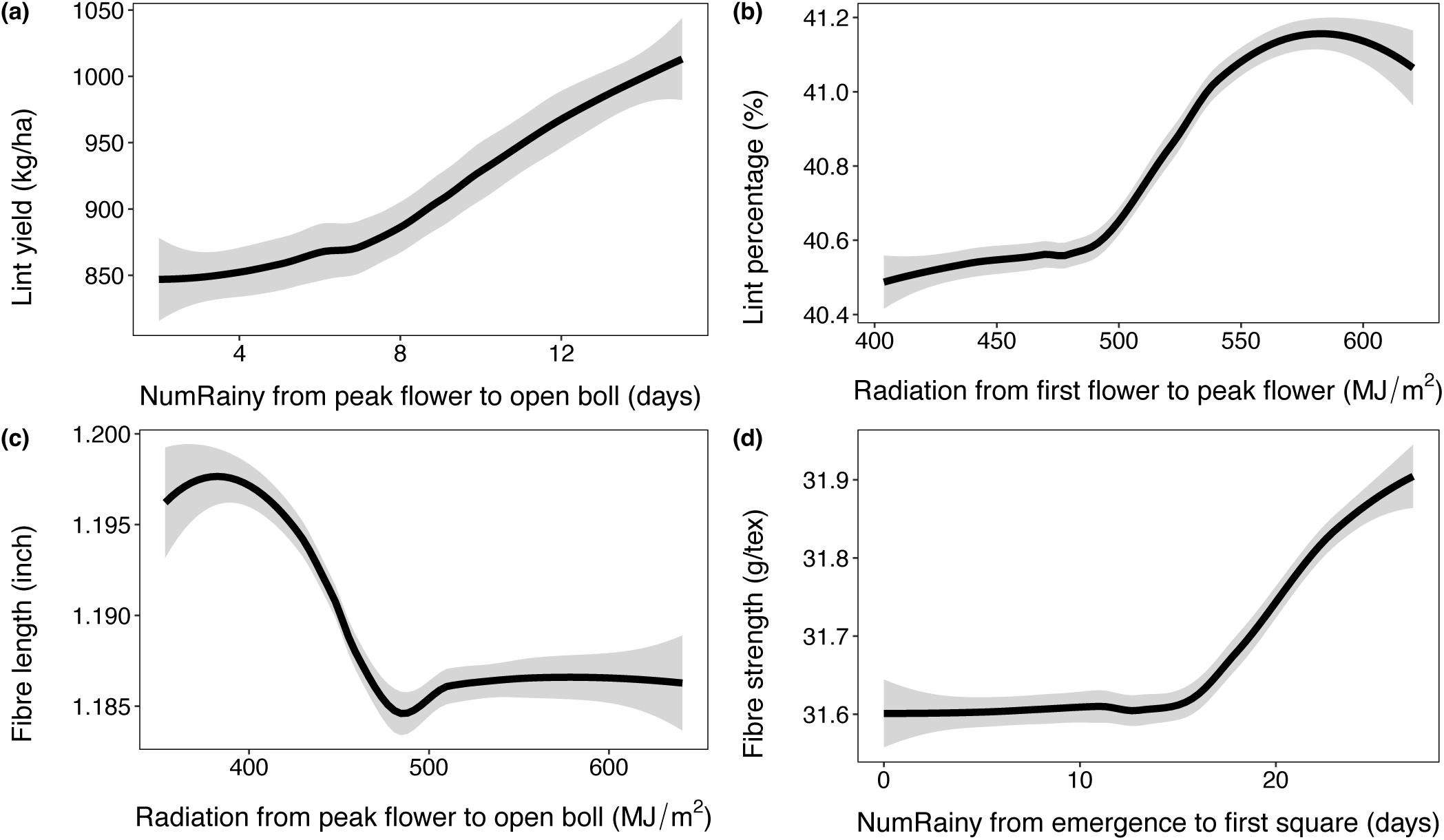
The non-linear relationship between four economically important fibre yield and fibre quality traits (lint yield (a), lint percentage (b), fibre length (c) and fibre strength (d)) in Australian rain-grown cotton and their most influential environmental covariate. The grey area denotes the 95% confidence interval. NumRainy represents the number of rainy days, radiation represents cumulative solar radiation.

Cumulative solar radiation was found to significantly influence LP and length-related fibre quality traits (EL, LEN, SFI, UNI). Increasing the cumulative solar radiation from 400 to 570 MJ/m^2^ from the first flower to peak flower period was associated with an increase in LP (**Figure 6b**). In contrast, LEN, a key quality indicator, decreased as solar radiation increased from 400 to approximately 500 MJ/m^2^ from peak flowering to open boll period (**Figure 6c**). Finally, mean VPD_max_ significantly influenced all fibre quality traits but none of fibre yield traits. The importance of VPD_max_ was highest for EL (from planting to emergence), MIC (from peak flower to open boll), and UNI (from open boll to maturity). In addition, the relationships between these VPD_max_-related ECs and the traits were all non-linear (Figure S8).

## 4. Discussion

### 4.1. Comparison between machine learning methods for variable selection and prediction

One of the aims of this study was to evaluate which machine learning and statistical methods provide optimal solutions for selecting the most influential ECs and predicting trait performance. We chose a wide range of methods to represent distinct modelling paradigms and assumptions and subsequently compared their performance. Given the complexity of environmental predictors and under-examined relationship with the response, no single method was assumed to provide an optimal solution for both inference and prediction.

#### 4.1.1 Differences in variable selection across methods and rationale for consensus

Substantial differences were observed in the number of influential ECs identified by different variable selection methods (**Table 4**). Classical frequentist SR consistently selected the largest number of influential ECs across traits, whereas the tree-based ensemble methods RF and BART were the most restrictive. Intermediate levels of sparsity were observed for BGAM, LMM and MLGL. These patterns primarily reflect intrinsic methodological properties rather than inconsistent biological signals.

SR is a sequential frequentist procedure that evaluates predictors conditionally based on incremental improvements in model fit. In the presence of correlated environmental predictors, this approach tends to retain multiple variables with overlapping information, leading to relatively inclusive EC sets. In contrast, RF and BART perform implicit variable selection through recursive partitioning and regularisation, with significance assessed using permutation-based tests. This procedure results in conservative selection that emphasises the strongest signals while excluding weaker or partially redundant effects. MLGL and the Bayesian approaches balance shrinkage and flexibility, producing more moderate EC subsets.

Given these method specific tendencies, relying on a single variable selection method would bias inference towards either overly inclusive or overly conservative EC sets. This explains why we opted to the consensus strategy, which prioritises robustness by retaining ECs consistently supported across multiple modelling frameworks and reduces sensitivity to artefacts associated with individual methods.

In addition, these method dependent differences are consistent with previous variable selection studies in environmental, genomic, and enviromic contexts (Campos et al., 2019; Montesinos-López et al., 2023). Frameworks such as *EnvRtype* emphasised environmental characterization and biological interpretation rather than formal variable selection under model uncertainty (Costa-Neto et al., 2021), while other genotype-by-environment (G×E) studies incorporated ECs through mixed models or random regressions into without explicitly targeting covariate selection (Monteverde et al., 2019; Tolhurst et al., 2022). Our consensus-based approach complements these studies by explicitly addressing uncertainty arising from method-specific variable selection before dissecting G×E interactions, a challenge increasingly recognised in integrative G×E analyses (Li et al., 2015).

#### 4.1.2 Predictive performance across methods

The purpose of determining an optimal prediction model was to better study the relationship between the influential ECs and fitted trait values. We used all ECs as predictors rather than only influential ECs from Step 2, as variables contributing to prediction accuracy are not necessarily those that are most statistically robust.

From our results, RF achieved the lowest RMSE and MAE for seven of the eight traits analysed, with BART showing comparable performance across traits and marginally outperforming RF for SFI (**Figure 3** and Figure S3). Both ensemble methods (RF, BART) consistently outperformed classical frequentist (MLR), penalised (MLGL), Bayesian (LMM, BGAM), and deep learning (DNN) models. The strong predictive performance of RF is consistent with its capacity to model non-linear relationships and interactions among predictors without requiring explicit specification of functional forms. Fibre yield and fibre quality traits in cotton are influenced by stage-specific and interacting environmental effects (Yao et al., 2026; Souaibou et al., 2025), and RF is well suited to capturing such complexity.

These findings are consistent with previous research, confirming RF as a strong predictive tool. RF has been widely applied and repeatedly shown to outperform other ensemble and deep learning methods. For example, studies on cotton yield (Dhaliwal et al., 2022) and fibre quality (Souaibou et al., 2025) reported that RF outperformed gradient boosting (Friedman, 2001), artificial neural networks, and linear regularization approaches such as Lasso. For yield prediction of the non-cotton crops, RF outperformed the MLR consistently (Jeong et al., 2016) and other classical machine learning methods (Yang et al., 2024; Lionel et al., 2025).

We also assessed computational efficiency across methods using a high-performance cluster (HPC) at Commonwealth Scientific and Industrial Research Organisation. Using a single core in HPC, SR was the fastest among all variable selection methods studied, averaging 16.8s per trait. Whereas RF required about 5.2h, mainly due to hyperparameter tuning and permutation tests. BART was the slowest approach, taking 9.6h per trait. Although RF’s prediction step took only seconds once tuned, its tuning process dominated runtime. Parallelizing hyperparameter search could substantially reduce this cost (Figures S9-S10).

### 4.2. Influence of environmental covariates on fibre yield and fibre quality

#### 4.2.1. Fibre yield

Blanc et al. (2008) and Traore et al. (2013) noted that the number of dry days during the Malian rainy season were associated with reduced LY; this is consistent with the positive relationship observed between number of rainy days and LY in the present study. The importance of growth stage was also evident, with environmental conditions during the latter half of the growing season exerting the strongest influence on LY. Dhaliwal et al. (2022) observed that critical variables during the period from flowering to open boll were more important for LY than those during the period from first floral bud to flowering. Broughton and Conaty (2022) also noted a strong association between LY and VPD during mid-to-late reproductive stages. These findings correspond closely to the peak flowering to maturity period identified as important in the present study.

LP in the present study was primarily influenced by cumulative solar radiation and daily minimum temperature from first flower to peak flower. Pettigrew (2001) highlighted the role of solar energy in enhancing photosynthetic carbon assimilation and promoting assimilate allocation to lint. Thenveettil et al. (2025) studied the impact of temperature and found that high night temperature was associated with a sharp reduction in LP during flowering period. These results support the observation in the present study that LP showed a positive response to increasing radiation and decreasing daily minimum temperature during early flowering.

Importantly, both LY and LP were governed by environmental variations within certain phenological windows, reinforcing the broader finding that fibre yield components are more controlled by stage-specific environmental drives than by seasonal averages (Reddy et al., 1992; Pettigrew 2001).

#### 4.2.2. Fibre quality

The present analysis demonstrates that environmental impacts on fibre quality in Australian rain-grown cotton are highly trait and stage specific, strongly non-linear, and frequently dominated by multiple environmental factors rather than by a single meteorological variable. Three ECs (daily maximum temperature from emergence to the first square, solar radiation from peak flowering to the open boll, and the number of cold days from maturity to harvest stage) were identified in the present study as the most influential ECs for LEN (**Figure 5**). This pattern highlights that LEN is influenced by both energy balance and thermal conditions across multiple developmental windows. The negative impact of solar radiation suggests that excessive radiation may increase evaporative demand and reduce cell turgor, thereby constraining length development, which is consistent with the work of Cosgrove (2005) and Dhindsa et al. (1975). The importance of temperature during early and late reproductive stages indicates that LEN may be partly shaped by carry-over effects from early reproductive development and by structural variability induced during fibre maturation.

SFI is the proportion of short fibres less than 12.7 mm in a sample and was highly significantly correlated with LEN in the present study (*r* = -0.5, *P* < 2.2 × 10^−16^). We therefore expected these two traits to share some influential environmental factors. This was supported by the present study, in which solar radiation was also identified as the most influential environmental factor to SFI, although the influential growth stage occurred slightly earlier than that for LEN. The present results also suggest that SFI was affected by the greatest number of influential environmental factors across almost all growth stages, compared with other quality traits analysed. This is not surprising, as SFI is a composite trait that integrate multiple aspects of fibre development and is therefore cumulatively affected by irregular growth, structural weaknesses and mechanical damage (Reddy et al., 1999; Bradow and Davidonis, 2000; Hequet et al., 2006).

In the present study, both STR and UNI were governed by the number of rainy days or rainfall across various growth stages, as well as mean VPD_max_ during the mid-to-late reproductive stage. These results align with irrigation studies showing the beneficial effects of cooler or wetter conditions on STR and UNI (Reddy et al., 1999; de Araujo et al., 2022; Schaefer et al., 2018). In variance component analyses, several studies have reported that the genetic variation contributes most strongly to STR compared with other fibre quality traits, explaining approximately 60-80% of the variance (Raper et al., 2019; Li et al., 2022; Virk et al., 2023). This implies that the remaining 20-40% of the variance is explained by non-genetic factors. This corresponds closely to the results of the present study, in which around 20% of the variance in STR was explained by the environmental factors.

Bange et al. (2010) and Bange et al. (2021) demonstrated that mean temperature during the fibre thickening phase, which overlaps with the periods from peak flower to open boll, is positively associated with increased MIC. Irrigation deficit studies have also identified this period of cotton development as important for MIC (de Araujo et al., 2022; Pinnamaneni et al., 2021; Schaefer et al., 2018). Wang et al. (2017) reported that the waterlogging during the flowering stage negatively affected the MIC due to limited photosynthesis and carbohydrate generated during fibre thickening phase. These studies agree with the present study in which appropriate VPD_max_ from peak flowering to open boll period was positively associated with MIC. Beyond that, present results also indicate a negative association between extreme high VPD_max_ and MIC (Figure S8). Together, these findings indicate that optimal, rather than maximal, atmospheric demand favours fibre maturity/fineness.

Fibre elongation occurs during the first two or three weeks following anthesis (Stiff et al., 2012; Bai et al., 2024), and environmental sensitivity has therefore often been assumed to coincide with early reproductive stages rather than plant establishment. In contrast, the present results indicate that EL was most affected by the environmental conditions during both the early and late growing season, specifically from planting to emergence and from maturity to harvest. These findings suggest that early environmental stress may influence subsequent EL indirectly, potentially through legacy effects on canopy development, assimilate supply, or sink establishment (Bange et al., 2004; Kerns et al., 2016). Low temperatures during the maturity-to-harvest period may alter fibre drying, maturation, or the physical condition of exposed lint, thereby affecting fibre extensibility measured at maturity, rather than directly regulating fibre-cell elongation itself (Reddy et al., 1999; Bradow and Davidonis, 2000; Kim and Triplett, 2001). Overall, although the underlying biological mechanism remains unclear, this finding represents a novel hypothesis generated by data-driven modelling and warrants targeted physiological investigation.

### 4.3. Future work

This study provides valuable insights into the environmental drivers of cotton yield and fibre quality traits, while also highlighting several opportunities for future work to further enhance understanding and predictive performance.

First, the set of the ECs analysed, although focused and informative, represents a subset of potentially influential covariates. We examined ten environmental factors in this study, while other factors such as soil moisture, wind speed and atmospheric carbon dioxide have been identified as important in previous studies (Reddy and Zhao, 2005; Wu et al., 2022; Souaibou et al., 2025). Once consistent, long-term datasets become available across year-location combinations, the inclusion of such variables in future analyses offer a promising avenue to further improve model performance. Expanding the range of ECs may increase the proportion of trait variance explained and help clarify relationships for traits such as elongation (EL), where current evidence is less strongly aligned with existing biological or literature support. Moreover, incorporating a broader environmental context may strengthen trait prediction and improve representation of genotype by environment interactions. Similarly, integrating management practices and their interactions represents an important next step that could further enhance predictive accuracy.

In this study plant growth stages were defined using thermal time, which provides a practical and widely adopted proxy in the absence of detailed phenological observations such as flowering or boll opening dates. While direct phenology measurements would be ideal, thermal time has been shown to effectively capture cotton development and remains an appropriate and robust approach when historical observations are limited (Bange et al., 2022 and Lin et al., 2023). Future work incorporating observed phenology, where available, would enable refinement of growth-stage classification and may further strengthen biological interpretation.

In addition, correlations among ECs from present study were anticipated given the interconnected nature of environmental processes. The variable selection methods applied here reduce redundancy and ensure robust modelling; however, future studies could explore methods that more explicitly account for collinearity to further disentangle independent effects. Accordingly, the associations identified in this study should be viewed as influential and biologically informative (instead of independently causal), providing a strong foundation for deeper investigation.

Second, while aggregation of environmental variables within growth stages provides clear biological interpretability aligned with crop phenology, complementary approaches based on time-series modelling offer additional opportunities. In current analyses, functional regression applied to daily environmental data confirmed the strong predictive performance of RF models (Figures S5-S6), with accuracy comparable to growth-stage-based approaches (Table S2). Although time-series models can capture complex temporal dynamics, their outputs can be less readily interpretable in a biological context. Future methodological developments aimed at improving interpretability of time-resolved models could help integrate the strengths of both approaches, enabling better identification of temporally specific stress responses.

Third, modelling fibre traits across a common set of phenological growth stages enables consistency and comparability; however, different fibre traits are known to be governed by partially distinct biological processes that may occur at different developmental periods (Haigler et al., 2012; Jan et al., 2022; Prasad et al., 2022). This suggests an opportunity for future studies to develop trait-specific growth-stage definitions. Achieving this will require larger, high-resolution datasets that allow independent estimation of optimal developmental windows for each trait. As such datasets become available, they may support more targeted and biologically precise modelling frameworks.

Finally, this study is based on data from inland cotton-growing regions of northern New South Wales and southern Queensland, providing a strong regional foundation for analysis. Expanding this framework to include a broader range of environments such as additional rain-grown systems within Australia or globally, represents an important direction for future research. Increased environmental diversity would enhance the ability to capture phenotypic variation, strengthen the detection of key environmental drivers, and further improve model generalisability. While direct extrapolation of results to regions with substantially different conditions should be approached with caution, the workflow developed in this study that links growth stages with influential environmental factors provides a transferable and scalable framework that can be readily applied to other cotton-growing regions and plant breeding programs.

## 5. Conclusions

Rain-grown cotton production represents a promising avenue for expansion of the Australian cotton industry. Understanding how the environment influences rain-grown cotton productivity is important for several reasons. Importantly, assessing the presence and magnitude of G×E effects for key fibre yield and fibre quality traits would likely improve the rate of genetic gain in rain-grown cotton breeding. Rather than incorporating an arbitrary set of ECs into the G×E analysis, the key ECs identified in the present study provide a solid foundation for future analysis. Furthermore, the statistical and machine learning methods used here could be extended to other crops, facilitating the translation of this research into broader breeding contexts aimed at improving crop performance.

## Credit authorship contribution statement

**Warren Conaty:** Conceptualisation, Project administration, Funding acquisition, Writing - review and editing. **Zitong Li:** Conceptualisation, Methodology, Supervision, Writing - review and editing. **Iain Wilson:** Conceptualisation, Writing - review and editing. **Pierce Rafter:** Writing – original draft, review and editing. **Qian Feng:** Formal analysis, Methodology, Visualisation, Writing – original draft.

## Data availability

The data used in this study are available for download via the CSIRO Data Portal upon the acceptance of the manuscript.

## Acknowledgements

The authors are grateful to Dr. Warwick Stiller, Dr. Shiming Liu, Dr. Tim Allen, and Dr. Annelie Marquardt for assisting with data curation and providing insightful feedback on the aim and direction of this study.

## Abbreviations

BART: Bayesian Additive Regression Tree
BGAM: Bayesian Generalized Additive Model
DNN: Deep Neural Network
LMM: Linear Mixed Model
MCMC: Markov Chain Monte Carlo
MLGL: Multi-layer Group Least Absolute Shrinkage and Selection Operator
MLR: Multiple Linear Regression
RF: Random Forest
SR: Stepwise Regression
VPD: Vapour Pressure Deficit
VPD_max_: Vapour Pressure Deficit at maximum daily temperature

