## supplementary for "Identification of environmental factors and growth stages in the prediction of fibre yield and fibre quality traits in rain-grown cotton"

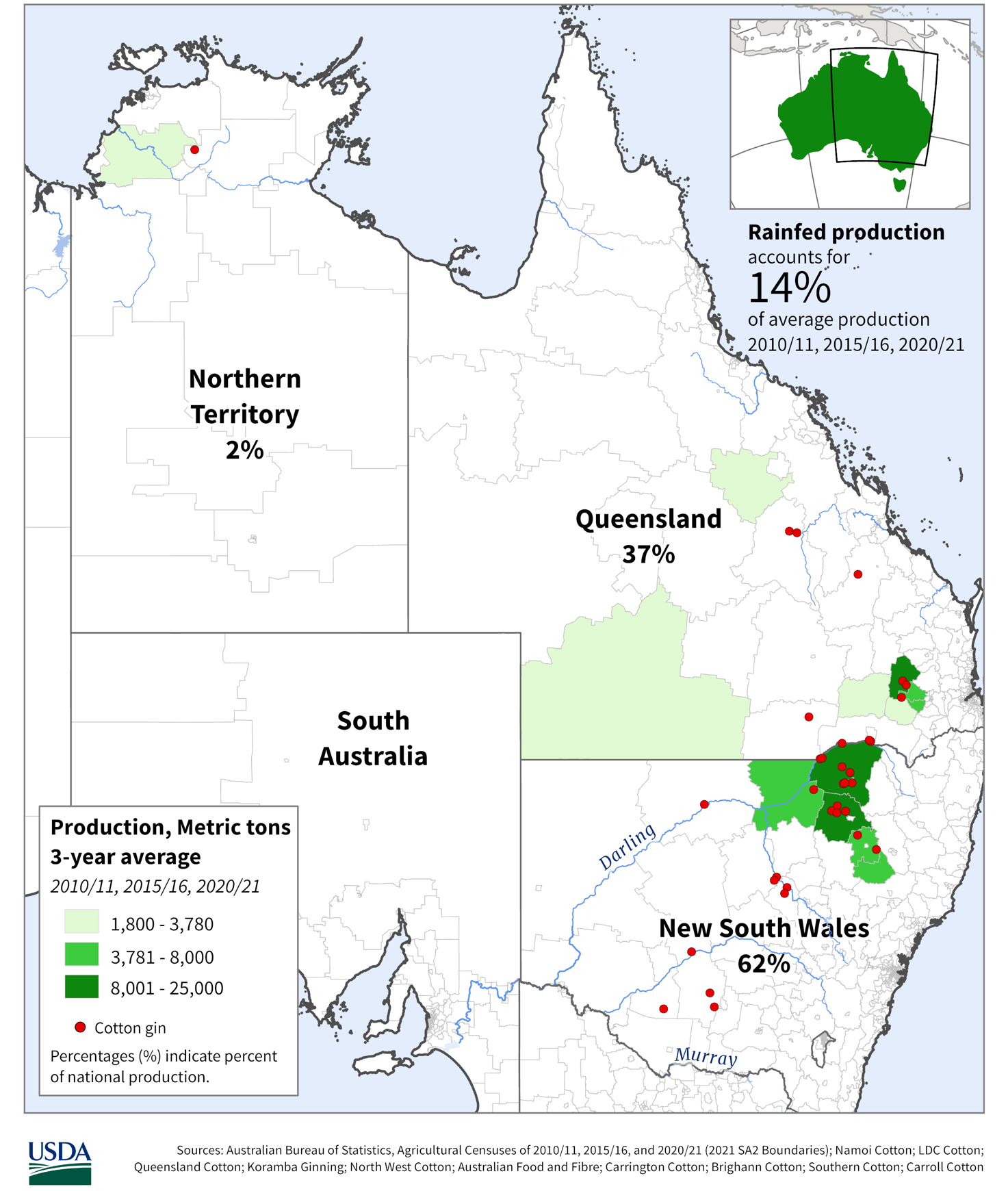

**Figure S1. Rainfed cotton production areas in Australia.** Adapted from “Australia Rainfed Cotton” [Map], U.S. Department of Agriculture, Foreign Agricultural Service, 2025 (<https://ipad.fas.usda.gov/rssiws/al/crop_production_maps/Australia/Australia_Rainfed_Cotton.png>).

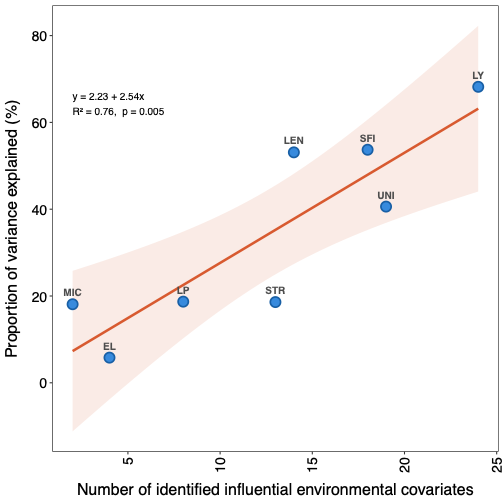

**Figure S2. Linear relationship between influential environmental covariates (ECs) and the proportion of variance explained across traits.** Each filled dot represents a trait. The proportion of variance explained was computed by the ECs identified from present study using a multiple linear regression. EL, LEN, LP, LY, MIC, SFI, STR, UNI represents for fibre elongation, fibre length, lint percentage, lint yield, micronaire, short fibre index, fibre strength and fibre uniformity separately.

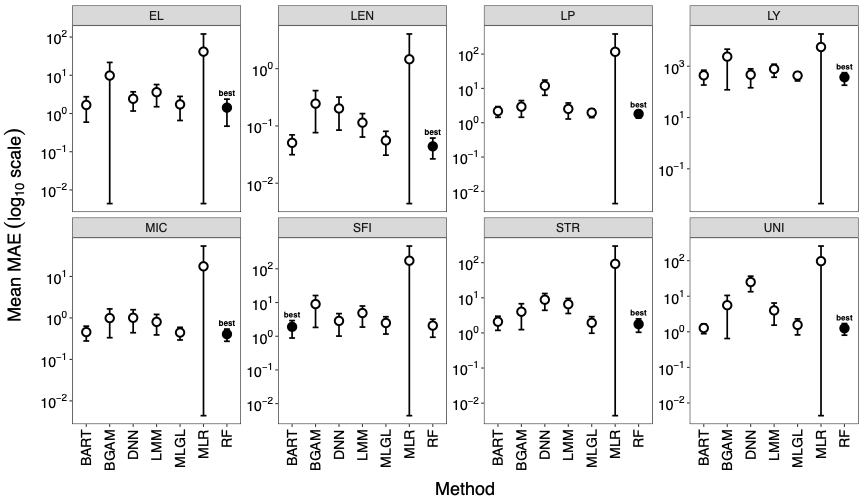

**Figure S3. Prediction accuracy measured by mean absolute error (MAE) of seven different methods for predicting fibre yield and fibre quality traits of Australian rain-grown cotton.** Each error bar refers to the Mean $\pm$ SD MAE across years, and filled circle denotes the best method (smallest MAE). BART, BGAM, DNN, LMM, MLGL, MLR and RF represent the Bayesian Additive Regression Tree, Bayesian Generalized Additive Model, Deep Neural Network, Linear Mixed Model, Multi-Layer Group Lasso, Multiple Linear Regression, Random Forest.

Supplementary text to

**B-spline in modelling time series environmental data**

In the present environmental data, seven factors had available daily records: Rainfall, RHmaxT, RHminT, Solar radiation, T.Max, T.Min, $\mathrm{VPD}_{\max}$. To model the changing trends with time, from the sow date to harvest date, of these environmental factors, we used B-spline function regression. This method works by combining simple polynomial pieces, with a knot vector specifying where the pieces connect. B-spline function regression is well suited for modelling time series data robustly against noise (Bak et al., 2025) and can handle observations with unevenly spaced time intervals, making it appropriate for our dataset.

In implementation, the raw time series (date sequences from the sow to harvest date) per year, location and environmental factor were normalised to the interval [0,1]. The break time points in the knot vector, except for the two endpoints, were defined based on the boundaries of seven growth stages. This resulted in eight break time points in total, $t\left( 0 \right), t\left( 1 \right),\cdots,t\left( 6 \right), t\left( 7 \right)$, where $t\left( 0 \right)=0, t\left( 7 \right)=1$. Each pair of adjacent break time points represented the duration of a growth stage. We constructed the polynomial pieces (basis functions) and implemented B-spline regression with “fda” R package (Ramsay et al., 2009; Ramsay 2025).

Finally, for each daily environmental factors, we obtained eight regression coefficients corresponding to the basis functions. In total these yielded 56 regression coefficients, forming a major component of the environment covariates (ECs) analysed. For the remaining three environmental factors (NumCold, NumHot, NumRainy), which do not have daily records, we counted their occurrence within each growth stage, resulting in an additional 21 ECs. In total, 77 ECs were used for subsequent variable selection and prediction.

**Table S1.** The number of identified influential environmental covariates (ECs) using 6 variable selection methods and consensus strategy for fibre yield and fibre quality traits of Australian rain-grown cotton. Input of each method was 77 ECs from our scenario 2. The consensus set was defined by the ECs identified $\geq4$ variable selection methods. See Abbreviations for the full name of each method.

| Trait | BART + Permutation | BGAM | LMM | MLGL | RF + Permutation | SR | Consensus |
| --- | --- | --- | --- | --- | --- | --- | --- |
| EL | 6 | 21 | 41 | 77 | 13 | 52 | 15 |
| LEN | 6 | 8 | 39 | 77 | 20 | 48 | 12 |
| LP | 2 | 8 | 37 | 38 | 17 | 46 | 2 |
| LY | 12 | 35 | 40 | 77 | 9 | 51 | 20 |
| MIC | 3 | 9 | 27 | 77 | 21 | 46 | 10 |
| SFI | 6 | 17 | 36 | 77 | 23 | 44 | 17 |
| STR | 1 | 14 | 35 | 38 | 15 | 50 | 6 |
| UNI | 8 | 13 | 28 | 77 | 18 | 50 | 15 |

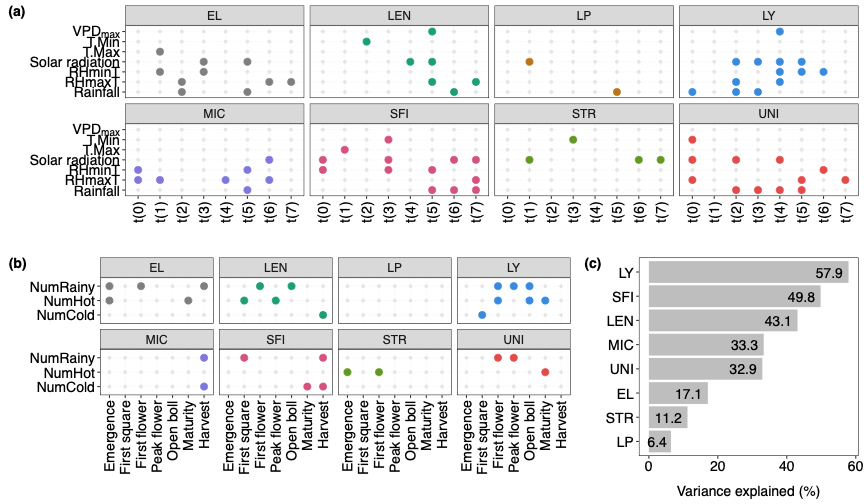

**Figure S4.** (a) (b) influential environmental covariates identified for fibre yield and fibre quality traits of Australian rain-grown cotton. Eight panels represent eight traits separately, the rows in each panel represent various environmental factors. The columns in (a) represents break time points across the knot vector, while the columns in (b) represents plant growth stages. If an environmental factor at a break timepoint/growth stage was found to influence a trait, a coloured dot is highlighted; (c) the proportion of variance explained by the identified influential environmental covariates. EL, LEN, LP, LY, MIC, SFI, STR and UNI stand for fibre elongation, length, percentage, yield, micronaire, short fibre index, strength and uniformity separately.

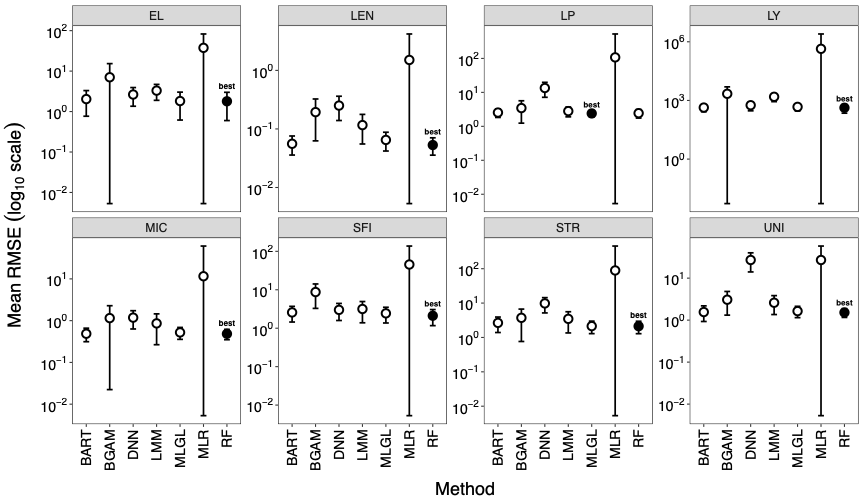

**Figure S5. Prediction accuracy measured by root mean square error (RMSE) of seven different methods for predicting fibre yield and fibre quality traits of Australian rain-grown cotton.** The predictors were 77 environmental covariates in scenario 2. Each error bar refers to the Mean $\pm$ SD RMSE across years, and filled circle denotes the best method (smallest RMSE). BART, BGAM, DNN, LMM, MLGL, MLR and RF represent the Bayesian Additive Regression Tree, Bayesian Generalized Additive Model, Deep Neural Network, Linear Mixed Model, Multi-Layer Group Lasso, Multiple Linear Regression, Random Forest.

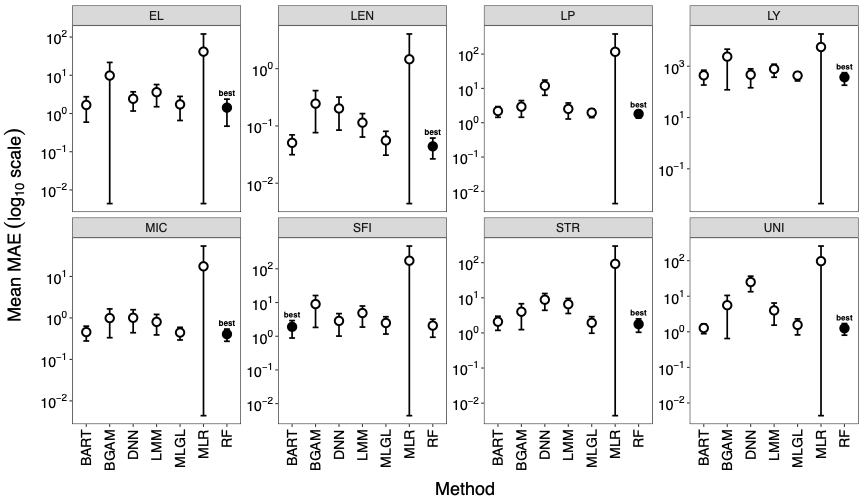

**Figure S6. Prediction accuracy measured by mean absolute error (MAE) of seven different methods for predicting fibre yield and fibre quality traits of Australian rain-grown cotton.** The predictors were 77 environmental covariates in scenario 2. Each error bar refers to the Mean $\pm$ SD MAE across years, and filled circle denotes the best method (smallest MAE). BART, BGAM, DNN, LMM, MLGL, MLR and RF represent the Bayesian Additive Regression Tree, Bayesian Generalized Additive Model, Deep Neural Network, Linear Mixed Model, Multi-Layer Group Lasso, Multiple Linear Regression, Random Forest.

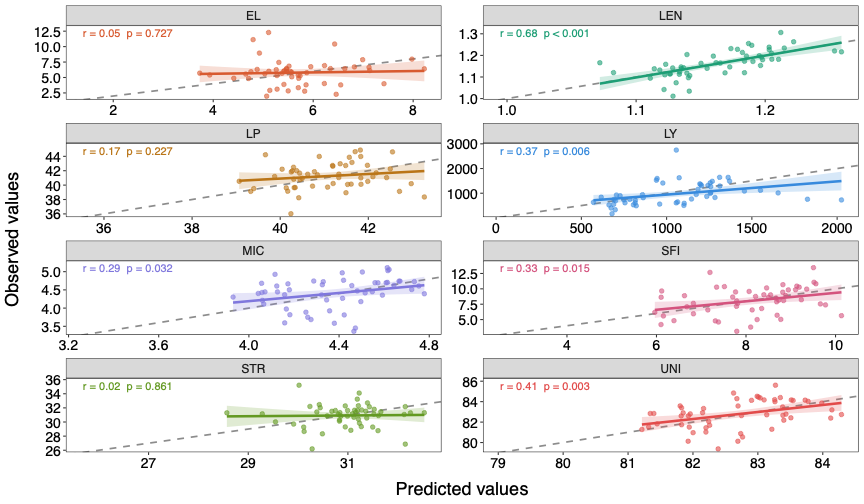

**Figure S7.** **Observed versus predicted mean trait values per year-location from the best-fit random forest model.** The predictors were 77 environmental covariates in scenario 2. Each panel represents a trait. Each dot in the panel represents a year-location combination. The dashed line is a 45% reference line, and the solid line is a linear relationship between the observed and predicted mean traits values. The shadow area denotes the 95% confidence interval. The correlation *r* between predicted and observed values and *p* value for the fitted regression relationship were reported in the top left of each panel.

**Table S2.** The average and standard deviation of prediction accuracy measures (RMSE and MAE) across years using the best-of-fit random forest model with and without daily environmental covariates modelled by the B-spline regression. EL, LEN, LP, LY, MIC, SFI, STR and UNI stand for fibre elongation, length, percentage, yield, micronaire, short fibre index, strength and uniformity separately.

|  | Mean RMSE (SD) | | Mean MAE (SD) | |
| --- | --- | --- | --- | --- |
|  | With | Without | With | Without |
| EL | 1.795 (1.196) | 1.676 (1.074) | 1.519 (1.039) | 1.422 (0.956) |
| LEN | 0.053 (0.017) | 0.054 (0.018) | 0.044 (0.017) | 0.044 (0.017) |
| LP | 2.439 (0.689) | 2.228 (0.487) | 2.005 (0.636) | 1.805 (0.441) |
| LY | 423.439 (199.993) | 435.975 (208.865) | 355.970 (164.273) | 371.739 (189.860) |
| MIC | 0.484 (0.134) | 0.481 (0.145) | 0.413 (0.125) | 0.406 (0.132) |
| SFI | 2.111 (0.945) | 2.348 (1.110) | 1.807 (0.942) | 2.072 (1.138) |
| STR | 2.124 (0.837) | 2.220 (0.859) | 1.679 (0.712) | 1.775 (0.733) |
| UNI | 1.520 (0.361) | 1.502 (0.484) | 1.265 (0.322) | 1.256 (0.452) |

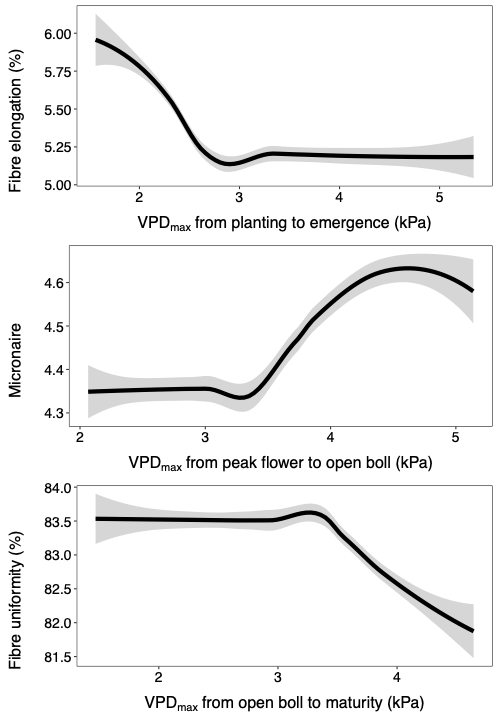

**Figure S8. The non-linear relationship between the fitted fibre quality traits (fibre elongation, micronaire, and fibre uniformity) in Australian rain-grown cotton and their top influential environmental covariate (**$\mathbf{VPD}_{\mathbf{max}}$ **at different growth stages)**. The grey area denotes the 95% confidence interval.

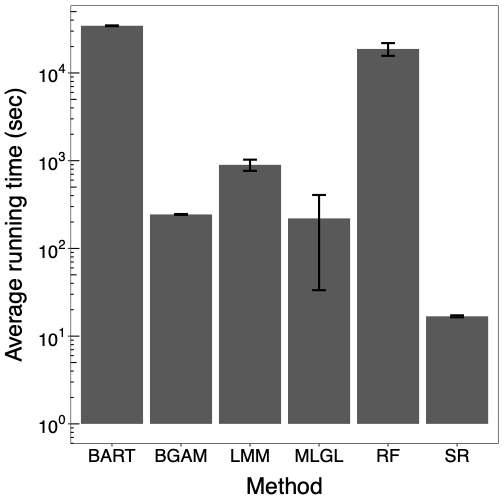

**Figure S9. The average running time across eight fibre yield and fibre quality traits of Australian rain-grown cotton with six variable selection methods.** The error bar represented the standard error of running time across traits. Since the running time varied a lot across methods, the y axis was log10 transformed for better visualisation.

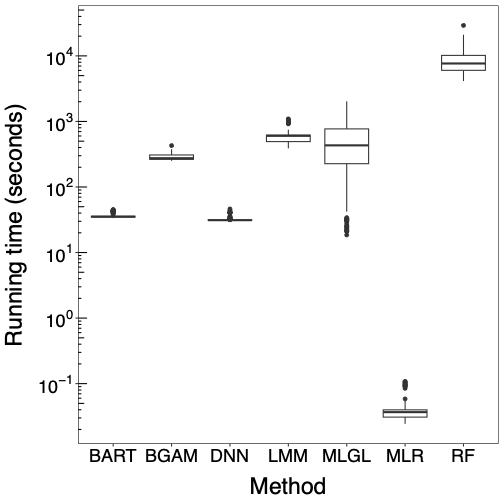

**Figure S10. The running time with seven prediction methods for eight fibre yield and fibre quality traits of Australian rain-grown cotton.** For each method used, we recorded the running time across years during leave-one-year-out cross-validation for all traits analysed and showed with a boxplot per prediction method. Some black horizontal lines appeared in this figure, note they were still boxplots just with narrow width. Since the running time varied a lot across methods, the y axis was log10 transformed for better visualisation.
